# Performance verification of human field of view occluders for light measurement and simulation

**DOI:** 10.64898/2026.08.04.742779

**Authors:** J. Mardaljevic, S.W. de Vries, J. van Duijnhoven

## Abstract

The measurement of light received at the cornea of the eye is a paramount consideration for the understanding of the relation between environmental illumination and the non-image-forming effects of light. The field of view (FOV) at the cornea is less than a full hemisphere, because it is partially occluded by human facial morphology. The International Commission on Illumination (CIE) has defined a standard model of human FOV. A suitably designed physical occluder attached to the sensor (of a light meter) has been proposed as a means of incorporating the effect of human FOV when taking measurements. Similarly, when using simulation to predict light received at the cornea, a geometrical description of the occluder at the eye point(s) can be added to the 3D model of the scene. The first occluder model proposed to represent CIE human FOV was enumerated in terms of: the CIE definition; the radius of the occluder; and, the radius of the light sensor disc. We present a simpler model based only on the CIE definition and the occluder radius. Both models were tested using a virtual goniophotometer. Various sensor response functions describing the spatial sensitivity across the sensor disc, including several we characterized through laboratory measurements, were included in the test. For all functions considered, the performance of the simpler occluder model was equivalent to or better than the model first proposed.

## 1 Introduction

Light affects various aspects of human health and well-being, including sleep, alertness, and mood. These effects are often referred to as the non-image-forming (NIF) effects of light and are largely attributed to the activation of intrinsically photosensitive retinal ganglion cells (ipRGCs) in the human eye (Dijk and Archer 2009; Lucas et al. 2014; Blume et al. 2019; Xiao et al. 2021). Consequently, assessing whether an environment’s lighting can support such NIF effects, for example, by evaluating it against recommendations (Brown et al. 2022), requires quantifying the (spectrally resolved) light a person receives at the eyes, often referred to as personal light exposure (PLE).

Using light meters, i.e. devices that measure incident visible radiation, two main approaches can be applied to quantify PLE. The first approach involves placing (typically laboratory-grade) light meters at locations representative of the eye position, for example oriented vertically at a height of 1.2 m in front of a desk. An estimate of an individual’s PLE can then be derived by combining these measurements with (approximations of) the time spent at each location. The same principle can also be applied using lighting simulations, in which PLE at representative positions is simulated rather than measured (e.g. see (Andersen et al. 2012; Mardaljevic et al. 2014; Danell et al. 2020; Alight and Jakubiec 2026)). The second approach involves subjects wearing a small light meter (dosimeter), preferably near the eyes (Spitschan et al. 2022; de Vries et al. 2025, 2026a). This approach is commonly used in field studies on NIF effects because it is relatively straightforward to apply across large participant samples and enables continuous monitoring in most environments to which subjects are exposed (Hartmeyer et al. 2023; van Duijnhoven et al. 2025).

Importantly, these approaches quantify light exposure at or near the eyes, whereas ipRGCs are activated by retinal light exposure. As such exposure cannot be measured directly (Spitschan et al. 2022), the next best proxy is the light reaching the human cornea (Zauner et al. 2023), where the field of view (FOV) is partially occluded by anatomical features such as the brow, nose, and eyelids (Sliney 1983, 2019), Figure 1a. Studies have shown that, depending on the illumination environment, measurements taken with an unoccluded (hemispherical) FOV can be substantially higher than those that account for FOV occlusion of the human eye (van Derlofske et al. 2000; Sliney 2019; Zauner et al. 2023; Broszio et al. 2025; de Vries et al. 2026b). Consequently, attaching a physical occluder (or hood) to light meters to mimic the human FOV has been proposed as a means to better estimate corneal light exposure (van Derlofske et al. 2000; International Commission on Illumination 2018; Zauner et al. 2023; Broszio et al. 2025).

**Figure 1:**
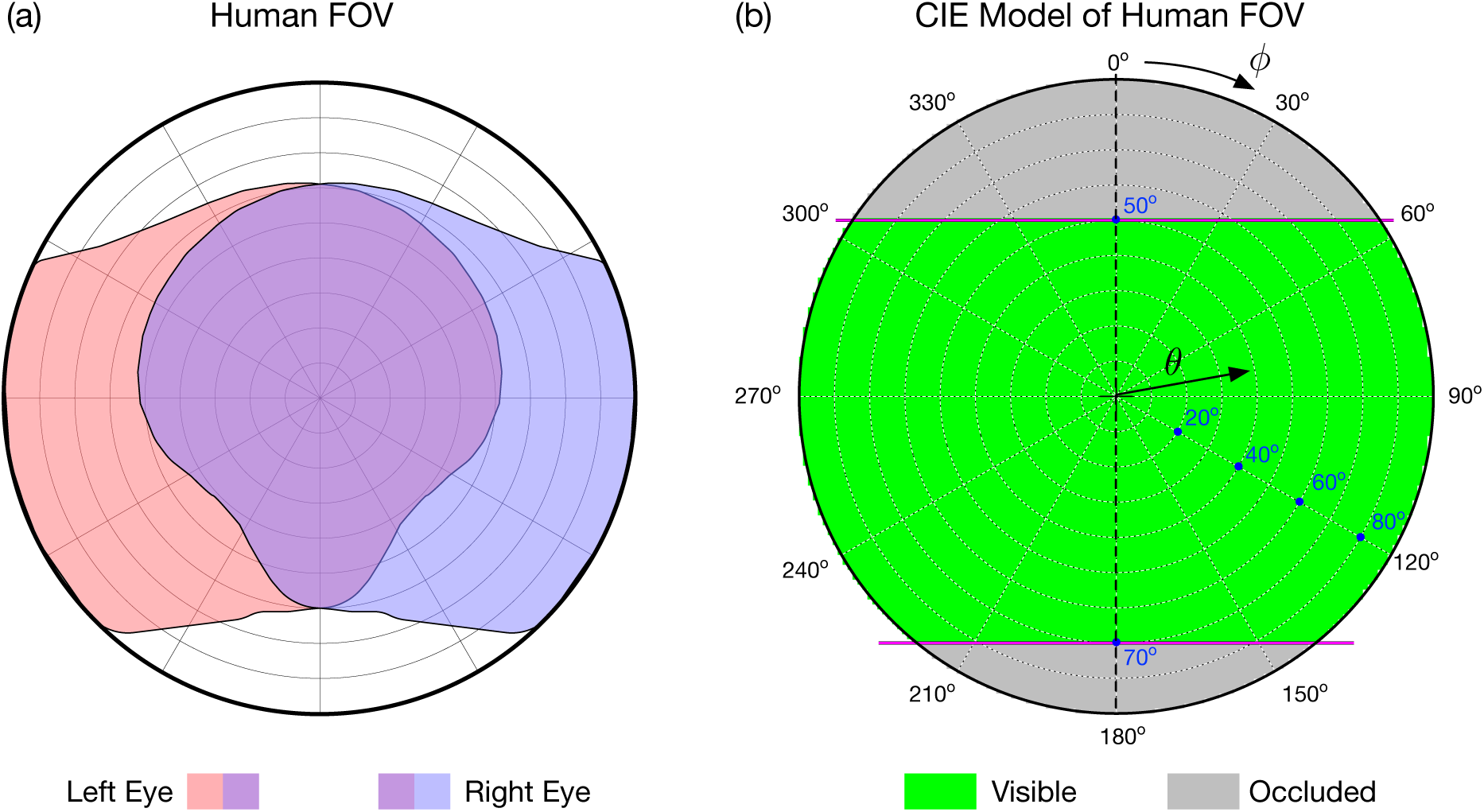
Human field of view (FOV) for left eye, right eye, and region of overlap (a) and CIE S 026:2018 definition (b) shown using equidistant angular fisheye projections. The coordinate system used in this article has *θ* as the angle of incidence to the view direction and *ϕ* as the clockwise angle around the view direction.

To facilitate the design of human FOV occluders, the International Commission on Illumination (CIE) specified standardized limits of the human FOV in CIE S 026:2018 (International Commission on Illumination 2018). Although left and right eyes have different occlusion profiles (Figure 1a), their aggregate effect forms the basis of this model. The CIE definition is graphically presented as an equidistant angular fisheye projection, centered on the view direction, which has horizontal and vertical view angles of 180*°*. There are two zones of occlusion, referred to as upper and lower, which are defined by two horizontal chords at 50*°* and 70*°* ‘above’ and ‘below’ the view center, respectively, Figure 1b. These values apply indoors; under brighter light (e.g. outdoors) the upper occlusion zone is larger. While this representation is generalized and thus does not capture individual anatomical variability, it provides a standardized method and enables comparison across studies.

The CIE’s FOV occlusion limits can only be exact for an infinitesimally small sensor. For a sensor of finite size, each position on the sensor disc — the outermost (diffusing) part of the sensor — views the occluder from a slightly different point, so the occlusion profile varies across the disc. How strongly this affects a reading (e.g. illuminance) depends on the sensor’s spatial responsivity, i.e. how its responsivity is distributed across the disc. The effect of an occluder therefore depends not only on the occluder’s shape, but also on the size and spatial responsivity of the sensor disc. To our knowledge, however, neither the performance of such occluders relative to the CIE definition has been tested photometrically, nor has the spatial responsivity of typical light meter sensor discs been characterized.

Here, we investigate how to design an occluder that best approximates the CIE FOV occlusion limits, accounting for both the size and the spatial responsivity of the sensor disc. To this end, we developed a virtual goniophotometer that computes partial-shading effects across the sensor disc, and characterized sensor responsivity profiles through laboratory measurements. We apply this approach to evaluate two recently proposed FOV occluders by Zauner et al. (2023) and Broszio et al. (2025), and of two occluders that we derived analytically from the CIE definition. We describe these occluders in the next section, followed by a brief background: first, on why partial occlusion of a light sensor has received little attention to date, and second, on goniophotometry.

### 1.1 Human FOV occluders

The occluders proposed by Zauner et al. (2023) and Broszio et al. (2025) are based on the CIE S 026:2018 definition. The first is a 3D-printed attachment for the JETI spectroradiometer (Figure 2a), with an occluder radius of approximately 18 mm, positioned around a sensor disc of radius 3.5 mm. The second is integrated into the 3D-printed casing of a custom dosimeter, the LIDO, which is designed to be attached to the arm of a pair of spectacles (Figure 2b). Its sensor disc has a radius of 5 mm and the occluder a radius of approximately 8 mm.

**Figure 2:**
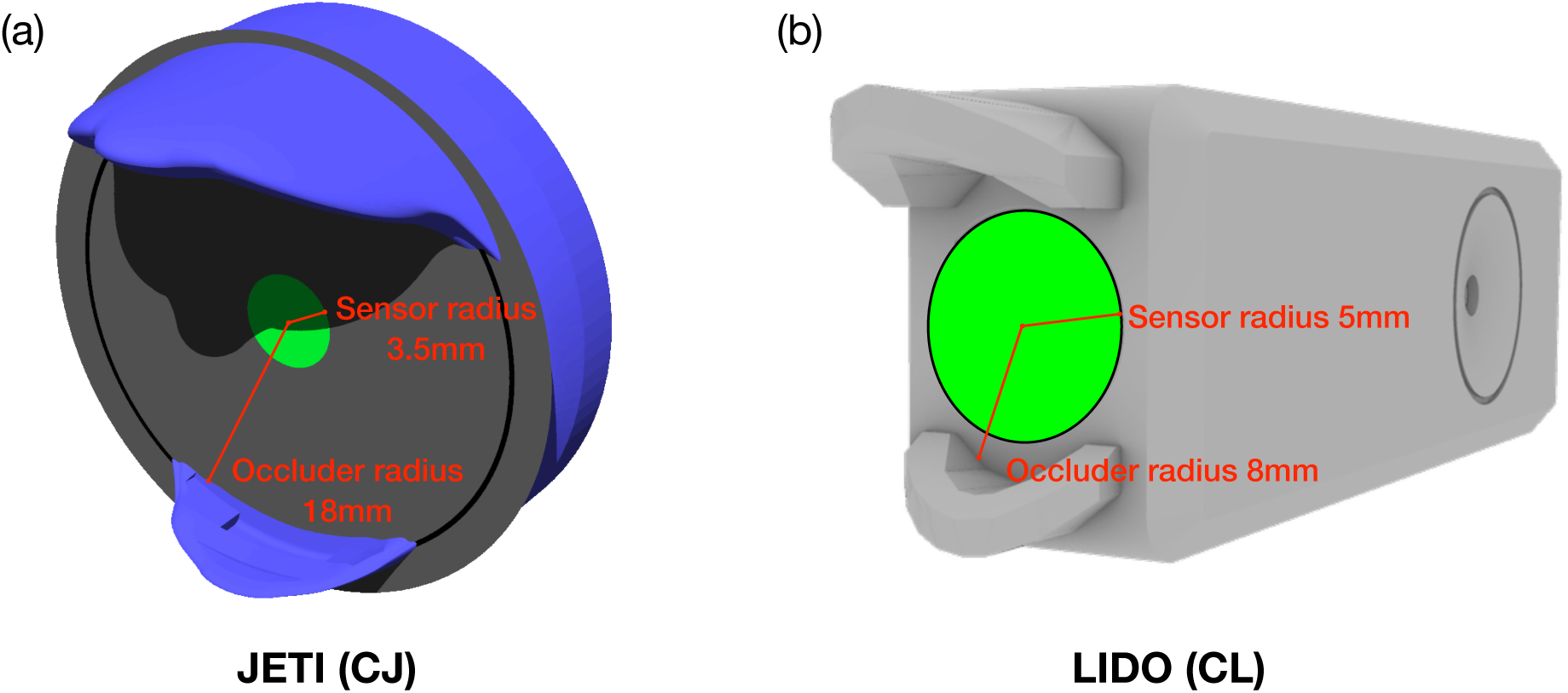
Renderings of the circumference-based JETI (CJ, a) and LIDO (CL, b) occluders. The JETI occluder is shown illuminated by a narrow light beam together with some ambient illumination, so that detail in the shadow regions is visible.

Both occluders were designed on geometrical principles that account for the area of the sensor disc: “The aperture [i.e. occluder] shape was then obtained by a line with the angle from a point on the edge of the diffuser [i.e. sensor disc] to the corresponding point on the opposite side of the aperture’s diameter.”(Zauner et al. 2023). The occluder shape can therefore be understood as ‘carved out’ from a construction line that follows the circumference of the sensor disc’s edge. We hence refer to these as circumference-based occluders, abbreviated CJ for the JETI occluder and CL for the LIDO occluder (Table 1). Importantly, the shape of such occluders differs with the radius of the sensor disc for which they were designed.

**Table 1:** Properties of the four investigated occluder designs. The circumference-based JETI (CJ) occluder was developed by Zauner et al. (2023) and the circumference-based LIDO (CL) occluder by Broszio et al. (2025). The effective radii of the CJ and CL occluders were estimated from the 3D models.

| Occluder design | Light meter | Abbr. | Sensor radius | Occluder radius | Occluder /<br>sensor radius |
| --- | --- | --- | --- | --- | --- |
| Circumference<br>based | JETI | CJ | 3.5 mm | ~18 mm | ~5.14 |
|  | LIDO | CL | 5 mm | ~8 mm | ~1.6 |
| Point based | JETI | PJ | 3.5 mm | 18 mm | 5.14 |
|  | LIDO | PL | 5 mm | 8 mm | 1.6 |

To illustrate how these occluders shade the sensor’s FOV, angular fisheye renderings of the view from three positions on the JETI’s sensor disc (from the center, the upper edge, and the lower edge) are shown in Figure 3. These parallax renderings were made using the *Radiance* lighting simulation system (Ward et al. 1998) and have a dimension of 1024×1024 pixels, with the sum of the solid angles of all pixels corresponding to 2*π* sr, with a precision of ±0.0001 sr. The part of the FOV obstructed by the occluder is shown in blue and the unobstructed view in gray. Each rendering also shows upper and lower chords (red) corresponding to the CIE definition, and a small inset graphic of the sensor disc (green) with a tiny red spot marking the viewpoint. From the center of the sensor disc (Figure 3a), the CJ occluder closely approximates the CIE definition, but not exactly. Small areas (that is, solid angles) that should be occluded are not, and vice versa – the upper and lower edges of the occluder ‘oscillate’ about the (red) chords. Considering next the occlusion from the point on the ‘upper’ edge of the sensor disc (Figure 3b), the upper occluder now presents a larger solid angle. As per the definition for the CJ occluder, the corresponding cut-off at the opposite point on the lower occluder, marked with a small red circle, is exactly at 70*°* (Figure 3b). Similarly, for the point at the ‘lower’ edge of the sensor disc (Figure 3c), the rendering shows the upper-occluder cut-off, as per the definition, exactly at 50*°*.

**Figure 3:**
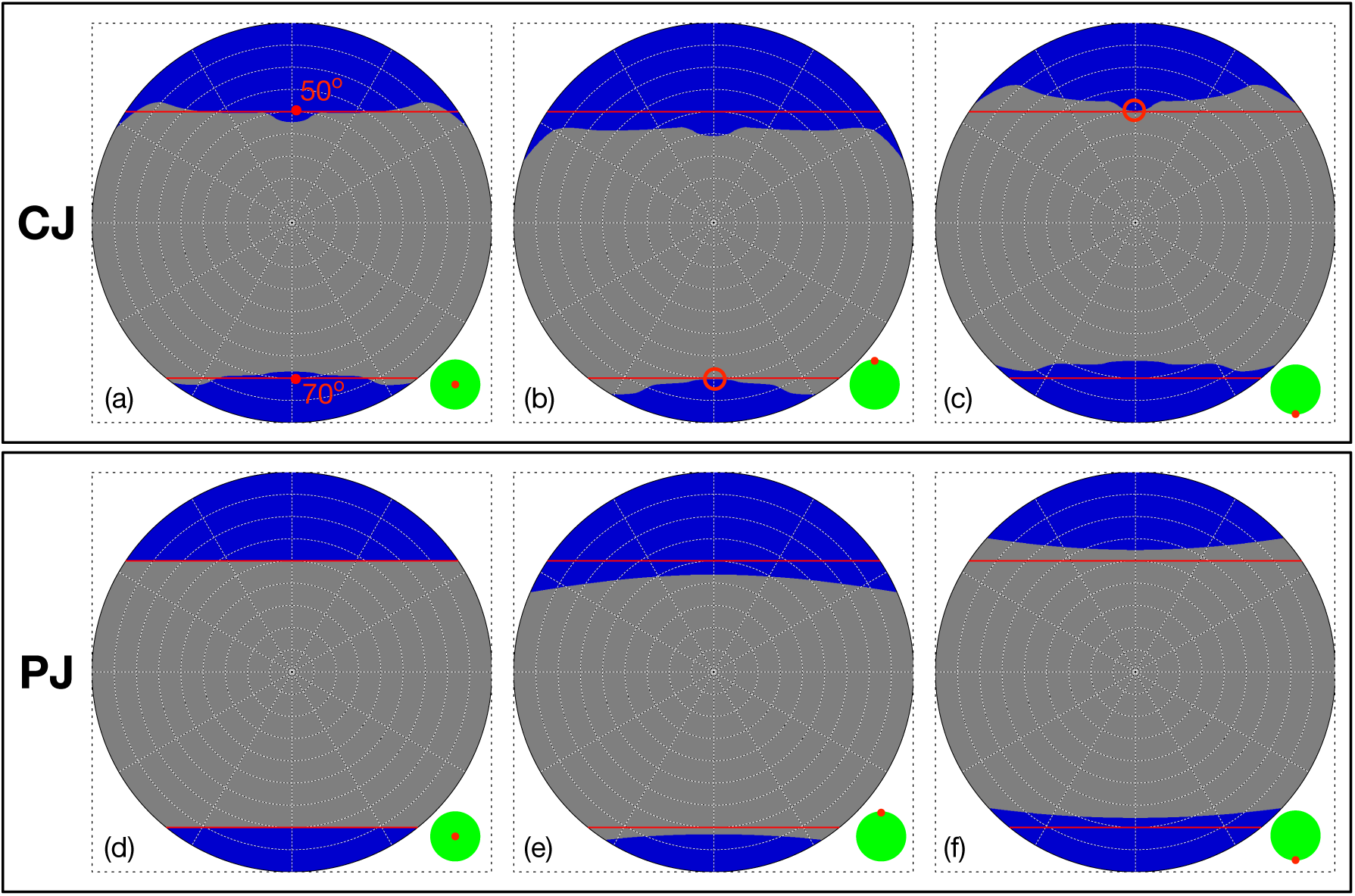
Angular fisheye renderings of the circumference-based JETI occluder (CJ) from the center of the sensor disc (a), and shifted by 3.5 mm (the sensor radius) ‘up’ (b) and ‘down’ (c). The same sequence of renderings for the point-based occluder (PJ) are shown in (d), (e) and (f). Blue pixels indicate obstruction by the occluder; gray pixels show the unobstructed view. Red chords mark the upper and lower bounds of the CIE S 026:2018 definition. The inset shows the sensor disc (green) with the rendering viewpoint marked in red.

The observations above motivated us to also consider a simpler occluder design, in which the shape is derived solely with reference to the center point of the sensor disc; the size (circumference) of the disc therefore is irrelevant to determining the occluder shape. Because the shape is fixed in this way, only its scale needs to be specified, which we define in terms of the occluder radius in the plane of the sensor disc. We refer to this ‘simple’ occluder design as a point-based model, and to the point-based model for the JETI device as the PJ occluder. The PJ occluder has the same radius as the CJ occluder (Table 1).

Point-based occluder models are facet models obtained from an exact analytical derivation of the CIE graphical representation. The PJ occluder model comprised 100 (four-vertex) facets each for the upper and lower parts, sufficient to very closely approximate the exact (infinitely narrow facet) representation. A rendering is shown in Figure 4a, and the adjacent sub-figure (4b) superimposes the mid-sections of the CJ and PJ occluders.

**Figure 4:**
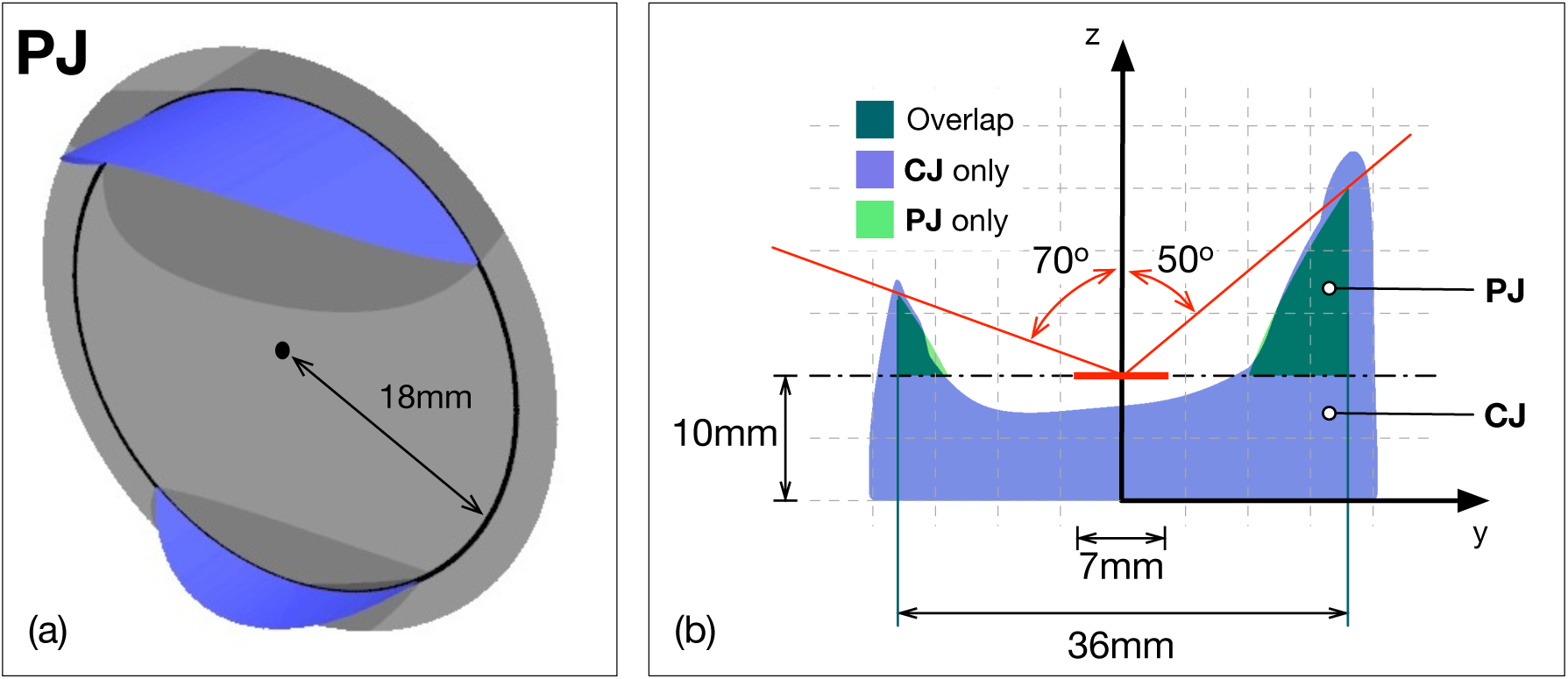
Rendering of the point-based JETI occluder (PJ, blue) with 18 mm radius (a), and mid-section of the PJ and circumference-based JETI (CJ) occluders superposed (b). occluder – and hence the more closely the CJ design approximates the PJ formulation.

The renderings of the view from the three positions on the JETI’s sensor disc are now repeated using the PJ occluder, shown in sub-figures (d), (e) and (f) in Figure 3. The first observation is that, viewed from the center of the sensor disc, the occlusion pattern exactly matches the CIE definition (d). From the shifted positions (e) and (f), the patterns show a parallax effect similar to that seen in sub-figures (b) and (c) for the CJ occluder. Here, however, the displacement from the CIE definition is more even with respect to the red chords. The CJ occluder has a largely similar shape to the PJ (Figure 4b) because the ratio of the occluder radius to the sensor disc radius is relatively large, 18 mm to 3.5 mm, or ~5.14 (Table 1). There is a parallel here with the so-called ‘five times rule’ in light measurement, which states that light from a luminaire should be measured at a distance equal or larger than five times the largest dimension of the emitting part of the light source (Moreno and Sun 2008). This ensures that the finite-sized source can be treated as a point source, yielding accurate, consistent measurements. Analogously, for the CJ occluder, the larger the ratio of occluder radius to sensor disc radius, the more closely the sensor disc approximates a point receptor at the center of the

The same renderings for viewpoints on the sensor disc of the LIDO device, considering the CL occluder and a point-based occluder with the same radius (referred to as the PL occluder, Table 1), are presented in Figure 5. As might be expected, the parallax effects are now much greater than those observed for the JETI (see Figure 3), since the ratio of the occluder radius to the sensor disc radius is considerably smaller at ~1.6 (Table 1). Considering first the CL occluder, from the center of the sensor disc the difference in the FOV from the CIE definition is quite marked: for the upper part, the occluder both encroaches on the FOV in the middle, but recedes too much at the edges, Figure 5, sub-figure (a). For the lower part, the occluder encroaches on the FOV by ~10*°* running almost parallel to the lower ‘chord’. In sub-figure (b), the parallax is such that almost the entire upper half of the FOV is obstructed. Similarly with the opposing point sub-figure (c), now the lower half of the FOV is obstructed to a marked degree. Turning to the PL occluder renderings, the first (sub-figure (d)) reproduces the ideal occlusion pattern (i.e. the same as in Figure 3d). For sub-figures (e) and (f), the parallax is conspicuous but less pronounced than for the equivalent CL occluder renderings, sub-figures (b) and (c) respectively. For each rendering (i.e. each viewpoint on the sensor disc) in Figures 3 and 5, the solid angles corresponding to the occluded and unoccluded FOV are available in Appendix A.1.

**Figure 5:**
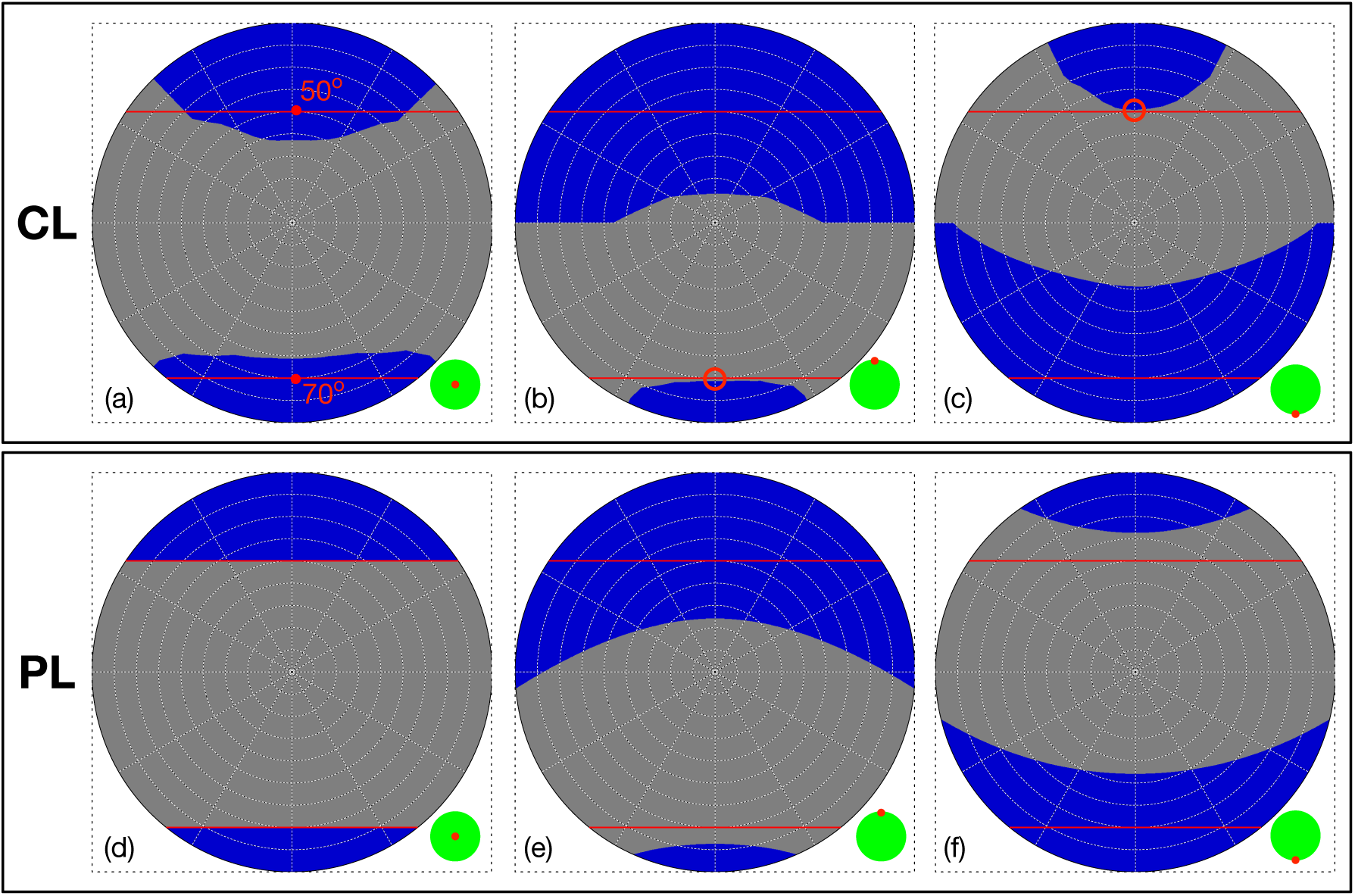
Angular fisheye renderings of the circumference-based LIDO occluder (CL) from the center of the sensor disc (a), and shifted by 5.0 mm (the sensor radius) ‘up’ (b) and ‘down’ (c). The same sequence of renderings for the point-based occluder (PL) are shown in (d), (e) and (f). Blue pixels indicate obstruction by the occluder; gray pixels show the unobstructed view. Red chords mark the upper and lower bounds of the CIE S 026:2018 definition. The inset shows the sensor disc (green) with the rendering viewpoint marked in red.

## 2 Background

### 2.1 Partial shading effects in illuminance measurements

The measurement of illuminance, or of related metrics based on other spectral weighting, with commercially available (calibrated) light meters is generally taken to be fairly straightforward. Although the manuals that accompany such devices can be extensive, the parts relating to the actual taking of measurements are usually brief. The technical specification for any particular device will often refer to a national standard, e.g. BS 667:2005 Illuminance meters – Requirements and test methods (British Standards Institute 2005). In terms of actual usage, the only guidance regarding occlusion of the light meter’s sensor is to avoid it, e.g. ‘*When performing measurement, take care not to allow the shadow or reflection of the operator to enter the receptor window.’* (from the Konica Minolta T-10A/T-10MA instruction manual, 2024). Although rarely, if ever, explicitly stated in such instructions, it is generally assumed that the sensor head will be exposed to an essentially uniform field of illumination at the point of measurement. In other words, small displacements of the order of the size of the sensor head should not result in a significantly different illuminance measurement.

In the literature, studies mentioning the effects of partial occlusion of a light meter’s sensor are sparse. In the context of the benchmark validation of the *Radiance* lighting simulation system (Ward et al. 1998) carried out in the late 1990s, simulated internal daylight illuminance predictions under real skies were largely within ±10% of measurements (Mardaljevic 1995). Nevertheless, there was a distinct class of ‘outlier’ predictions that were due to a number of factors related to visibility of the source (i.e. sun and/or circumsolar region) from the photocell sensor (Mardaljevic 2001). One of those was the effect of the partial occlusion of the sensor by window frame bars under sunny conditions: the actual sensor had a diameter of ~1 cm and could therefore be partly occluded, whereas the simulated sensor was modeled as a point.

The possibly confounding effect that objects close to the sensor head could have on light measurements is more documented. For example, the European standard for lighting of work places (EN 12464-1:2021) considers the impact of reflected lighting from walls on (grid-based) uniformity calculations: ‘*To avoid high impact on uniformity from calculation points near the wall, a band next to the wall can be excluded from the calculation […].’*. Moreover, such effects were discussed by Mardaljevic et al. (2021) in the context of a conservation setting. In that study, it was argued that the light meters placed directly on furniture tops would record illuminances markedly different from those that would be measured at the same point in space if the furniture were not present (see Figure 14 in (Mardaljevic et al. 2021)). Additionally, the material of the furniture tops had a marked specular component of reflection. In this position, the sensor would receive a grazing incidence specular reflection of light from a facing window. Objects placed close to a light meter’s sensor create a ‘localized illumination microclimate’, particularly when specular reflections are present. Accordingly, such locations could be both unrepresentative of the prevailing illumination, and also unreliable data points to compare against predictions. The *Radiance* system has also been used to study near field illumination effects due to parallax in the context of sky simulator domes (SSDs) used with physical models for illuminance prediction (Mardaljevic 2002). With a daylight simulation, the sun and sky are effectively at an infinite distance. Thus, whatever the dimensions of any building or city modeled in the simulation, every part of it ‘sees’ the same luminance pattern across the sky vault. In contrast, with a physical model placed in a SSD, any non-uniform luminance pattern on the sky dome and/or any sun position will only be ‘correct’ from one reference point. Consequently, different parts of the physical model will not ‘see’ the same luminance pattern on the dome. The resulting error for illuminance measurements depends on the ratio of the radius of the dome to size of the physical model (Mardaljevic 2002). Thus, a fundamental limitation of all (physical) sky simulator domes was revealed using simulation. Despite the very different context of the SSD parallax evaluation, there are strong conceptual similarities with the study reported here.

To summarize, until recently, and apart from a few niche areas of lighting science, the partial occlusion of, or interaction with, objects close to illuminance sensors was generally considered a potentially confounding factor that could be easily avoided by good practice. Namely, the user should not introduce additional shading at the point of measurement, and generally avoid nearby objects that could create a spurious illumination microclimate different from the prevailing illumination field which is the object of the measurement. However, as noted above, that has changed with the requirement to modify a light meter’s FOV using an occluder to account for the human FOV.

### 2.2 Goniophotometry

The term goniophotometer was originally used to describe a device for measuring the angular output in the distribution of luminous flux from a light source (Bertenshaw 2020). The earliest measurements date from the late 1800s, and the measured directional output of commercial luminaires began to be included in product specifications since at least the 1950s. In 1986, the first version of the ‘IES Recommended Standard File Format for Electronic Transfer of Photometric Data’ was published (Illuminating Engineering Society 1990). The standard was necessary because, with accurate photometric data, computers could reliably determine work plane illumination across large areas lit by hundreds of luminaires much faster than the tabular methods used previously. A schematic of the type of goniophotometer used to measure the light output distribution of luminaires is shown in Figure 6a. The luminaire is positioned at the origin with the emitting side (shaded yellow) directed ‘upwards’ (in this illustration, the opposite sense to how a ceiling-mounted light would be directed). Either the photometer or the luminaire (or sometimes a combination of both) would be incrementally rotated/moved so that the full hemisphere of light output from the luminaire would be recorded at the desired angular resolution. For the majority of ceiling-mounted luminaires, this is sufficient. However, suspended luminaires which also have some light directed upwards to illuminate the ceiling will need to be measured across the full hemisphere.

**Figure 6:**
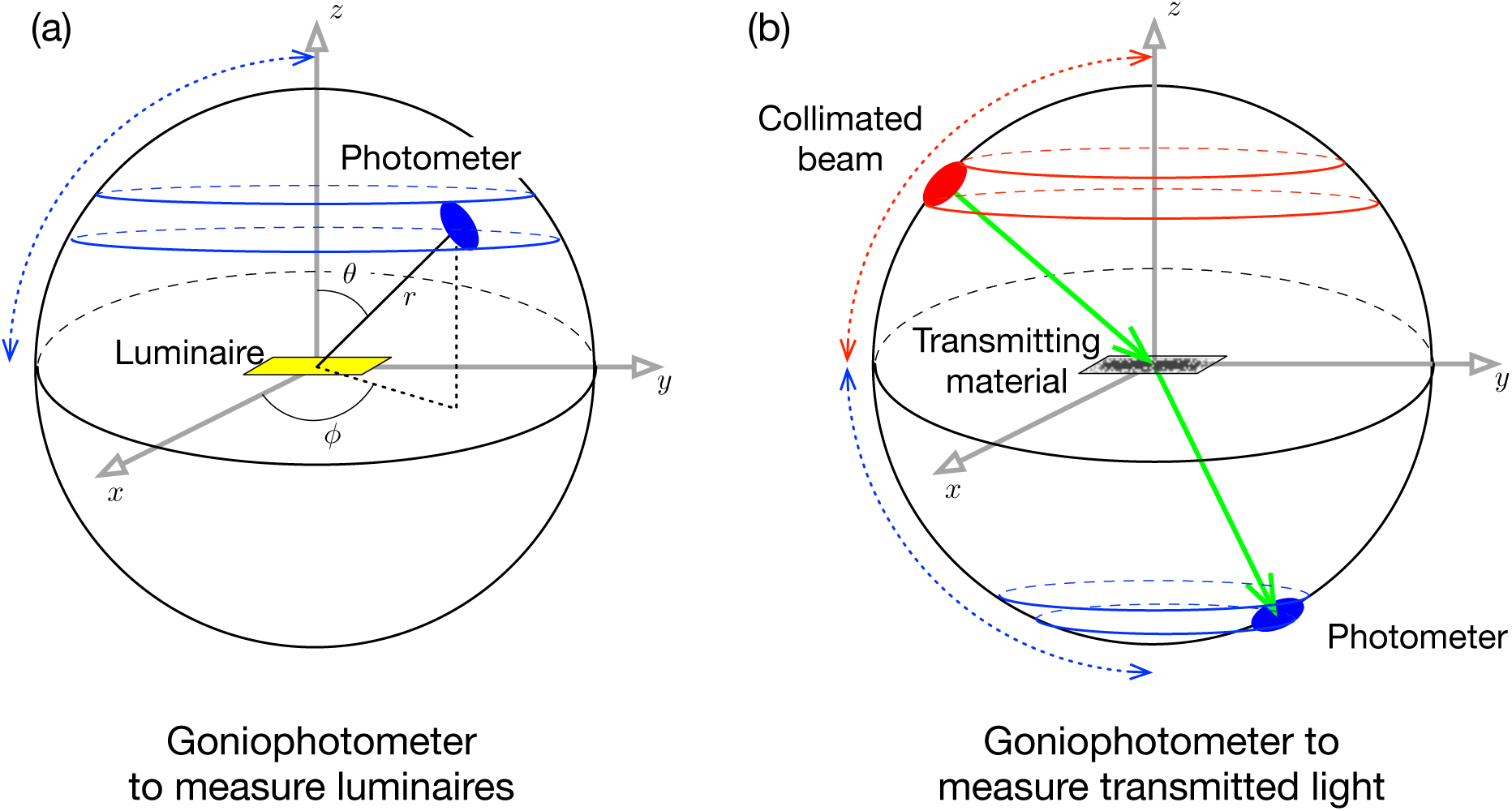
Schematic of the two most common types of goniophotometers, for measurement of luminaires (a) and transmitted light (b). The illustrations are annotated using the standard spherical coordinate system. The blue curves illustrate the ‘sweep’ paths of the photometer, and the red the ‘sweep’ paths of the collimated beam, i.e. to cover the respective hemispheres. Though actual movement will depend on the particular design of the goniophotometer. The coordinate systems used will vary depending on the nature of the particular application.

The next significant development in goniophotometry was the measurement of the light transmission properties of so-called complex fenestration materials (e.g. prismatic glazing) or devices (e.g. mirrored louvres). This required devices which could measure the light transmitted in all possible angles (i.e. a hemisphere) for all possible incident directions of light – also a hemisphere. A schematic is shown in Figure 6b. If the incident light were measured at *N* points distributed across the hemisphere, with the same distribution used for the transmitted light, then a total of *N* ^2^ measurements would need to be taken, i.e. *N* measurements of transmission for each of the *N* incident directions. The data that characterize the light transmission properties of a material or device for all possible incident directions of light are referred to as the bidirectional transmittance distribution function (BTDF) (Papamichael et al. 1988). To capture the distribution in reflected light from a material, the photometer would scan the hemisphere on the same side as the source. This characterization is known as the bidirectional reflectance distribution function (BRDF). The complete characterization containing both the ‘front’ reflected light and the ‘back’ transmitted light is called the bidirectional scattering distribution function (BSDF) (Bartell et al. 1981). Subsequent innovations used imaging techniques for more efficient data capture of the hemisphere of reflected and/or transmitted light (Ward 1992; Scartezzini et al. 1997).

A virtual goniophotometer is any simulation-based approach that reproduces the function of a physical goniophotometer. Simulation-based approaches were first considered in part because of the high cost, in terms of equipment, space, and time, of physical goniophotometry (Krishnaswamy et al. 2004). Since at least 2012, the *Radiance* lighting simulation system has included a specialized utility script called genBSDF to simulate BSDFs which can then be used in daylight simulations (Ward et al. 1998; Appelfeld et al. 2012). For example, genBSDF could predict the overall BSDF for a complex fenestration system comprising multiple material types, e.g. mirrored louvres, clear glazing, translucent sections, etc., provided the optical properties of each material type can be adequately described in the materials database.

## 3 Method

The method consists of three parts. First we describe the modeling of the sensor disc, then the virtual goniophotometer, and finally the measurement of response profiles across the sensor disc of a light meter.

### 3.1 Defining the light sensor disc

We describe the modeling of a flat sensor disc, of arbitrary size, as a collection of individual point sensors to which any assumed or measured response function of the sensitivity across the sensor disc can be applied. The sensor discs used in the JETI and LIDO devices have radii of 3.5 mm and 5 mm, respectively (Table 1). Each sensor disc was modeled as an ensemble of regularly spaced (receptor) points that lie within a defined circle, Figure 7. The coordinates of the points that represent the disc were determined using the image-based stencil method (Mardaljevic and Roy 2016). In short, the coordinates are extracted from a parallel projection *Radiance* image of the disc. Thus, the spacing between the points is determined primarily by the pixel dimensions of the image, rather than the size of the disc. For the work described in this article, 656 points were used to describe the sensor disc irrespective of the diameter of the disc. This number of points gives an accuracy of ~1% in the representation of the circular disc area by the yellow grid of square elements shown in Figure 7 (the circular area calculated from the radius to similar precision is 38.484 mm^2^).

**Figure 7:**
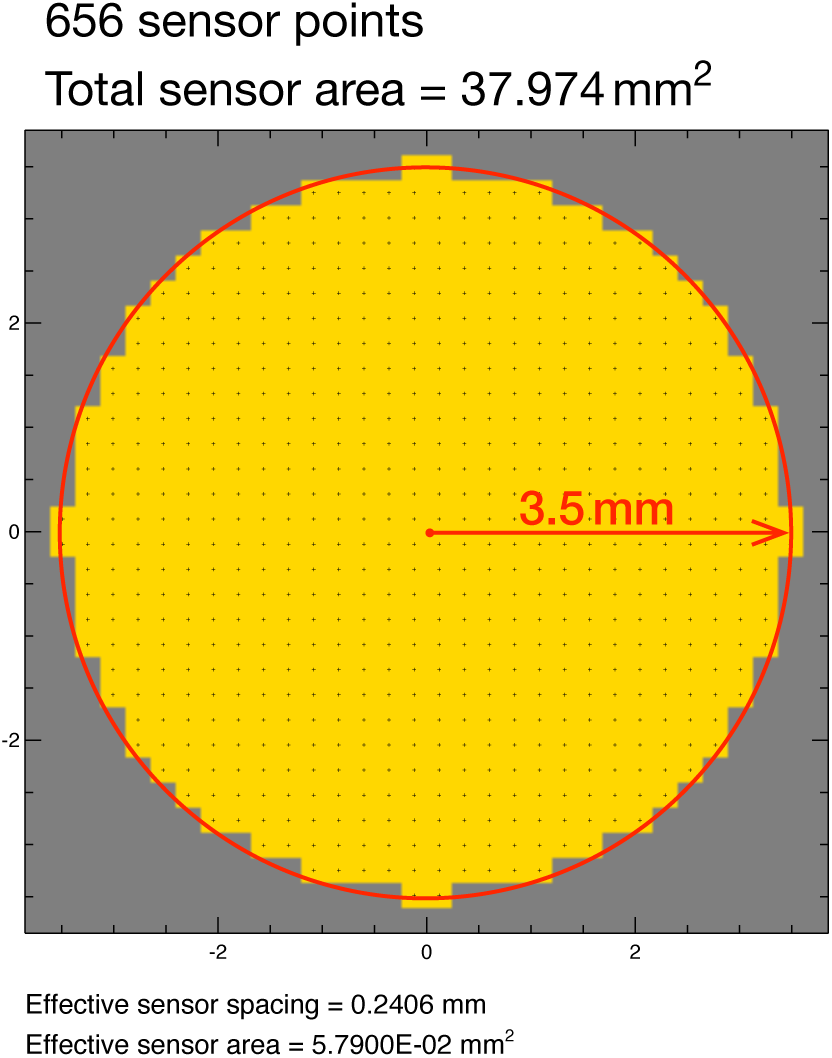
Illustration of sensor point pattern used to represent a 3.5 mm radius disc.

### 3.2 The occlusion virtual goniophotometer (OVG)

A virtual goniophotometer specifically tailored to characterize the effects of near-field occlusion on an illuminance sensor was developed using the *Radiance* lighting simulation system. Hereafter referred to as the occlusion virtual goniophotometer (OVG), it was designed on the basis of the equidistant angular fisheye projection so that the results would have a 1-to-1 correspondence with the graphical presentation of the CIE definition for human FOV (International Commission on Illumination 2018; Zauner et al. 2023). A schematic of the OVG is shown in Figure 8. A 2D Cartesian plane normalized to equal ranges of [-1,1] for both axes is shown superposed on an equidistant angular fisheye graticule (Figure 8a). The graticule shows circular lines at 10*°* intervals (from the view center) and ‘meridian’ lines at 30*°* increments. For this illustration, the Cartesian plane has been subdivided into a ‘coarse’ 16×16 grid. The center of each grid square which is inside the graticule (red dot) is used to generate the (*x, y, z*) vectors which define the angular coordinates of the associated goniophotometer point light source on the hemisphere. For the 16×16 grid, there are 208 points inside the graticule, and so there are 208 light sources distributed across the hemisphere. This distribution is illustrated in the rendering shown in Figure 8b. However in the OVG, each actual light is defined as a source solid angle, similar to how the sun is described in a *Radiance* scene file. Thus, the light source is effectively at infinity and, because a single ray is used to sample the source, the beam width is effectively zero, irrespective of the size of the solid angle. Physical goniophotometers limit the beam size using baffles and/or lenses to achieve a compromise between the signal strength and the angular size of the beam (Apian-Bennewitz and von der Hardt 1998).

**Figure 8:**
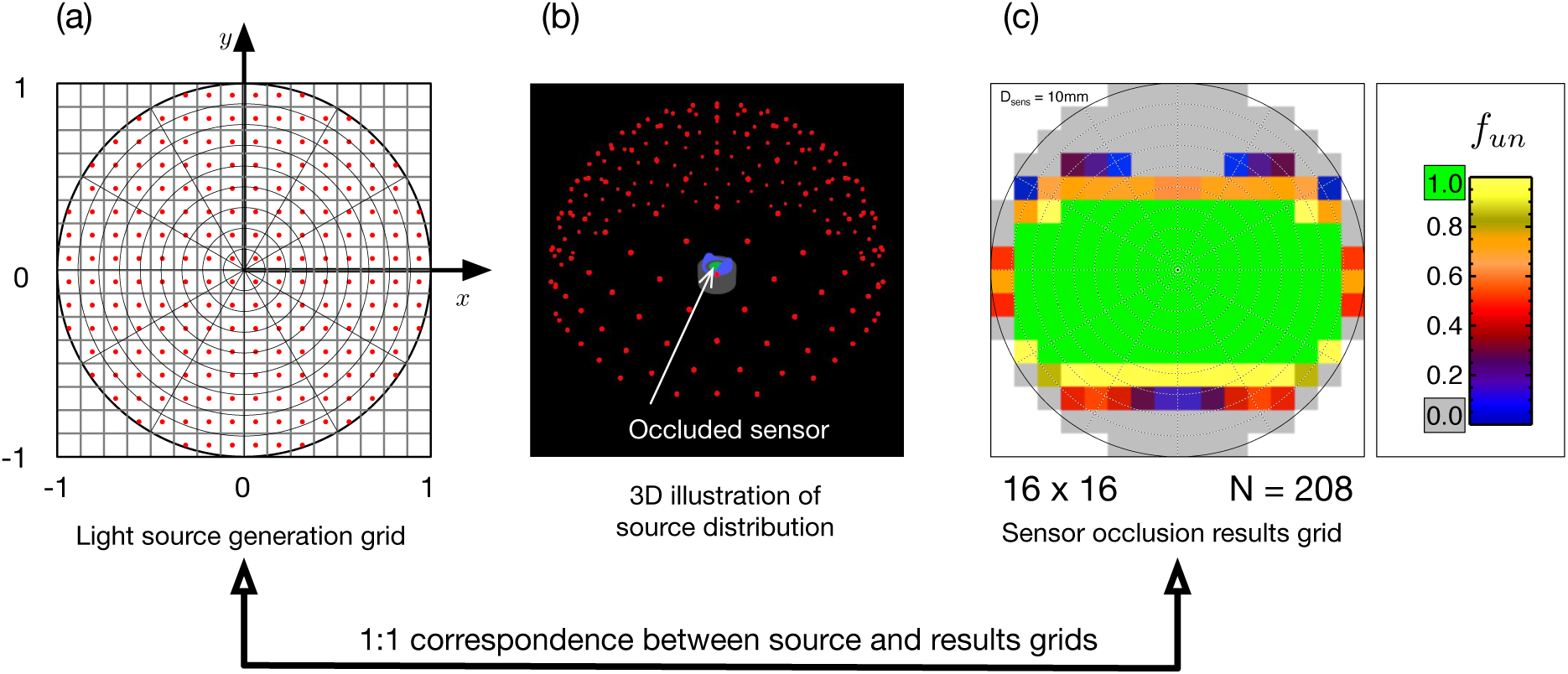
Schematic for the occlusion virtual goniophotometer (OVG) grid-based design with 1-to-1 mapping between source (a, b) and results grids (c). This illustration uses a low resolution 16×16 grid giving 208 light sources distributed across the hemisphere. An angular fisheye projection is used. The light sources are shown as red dots in (a, b) and the occluder and sensor are shown in blue and green, respectively, in (b).

The last part of the illustration (Figure 8c) shows an example of results for a 10 mm diameter sensor with an occluder. The circular sensor is modeled as a discretized grid of (receptor) points. For now, the purpose of the illustration is to show how each grid square is shaded to represent the fraction of the (circular) sensor area that received illumination from each of the 208 light sources. Each square therefore does not represent an actual location on the sensor disc; rather, it is linked to one light source, at an angular position indicated by the position of the square. The number of points on the sensor disc illuminated by that light source determines the unshaded fraction (*f_un_*). The squares shaded green indicate that, for the angular position at the square center, the light source illuminates every point of the sensor. Thus, the fraction of the sensor disc unshaded (*f_un_*) is one. For squares shaded gray, the unshaded fraction is zero; in other words, the occluder prevents light from sources located at the angular position of the (gray) square centers from reaching any of the points on the sensor disc. Partial shading (0 *< f_un_ <* 1) is indicated using the (linear) color scale shown in the legend. As can be seen (at this coarse resolution), there are a number of light sources around the unshaded (green) squares where partial occlusion of the sensor occurs.

The imposition of a 1-to-1 correspondence between the Cartesian projection of the OVG light sources and the computed shading on the sensor precludes any requirement for interpolation between the light source and results grids. This is important because, any applied interpolation (upscaling or downscaling) could inadvertently produce spurious patterns of partial shading where none occurred. The OVG was designed to accommodate (square) grids of arbitrary resolution, limited only by the computing resources available. The results from repeating the shading computation with progressive refinement of the grid are shown in Figure 9. Grids of 32×32, 64×64, and 128×128 were computed, giving, respectively, 812, 3,228, and 12,892 light sources distributed across the hemisphere. We determined from tests that the 128×128 grid gave sufficient resolution to reveal the significant performance differences between the circumference and point occluder types. The solid angle associated with each pixel in the 128×128 grid varies gradually from the periphery (~0.0004 sr) to the center (~0.0006 sr), Figure 10.

**Figure 9:**
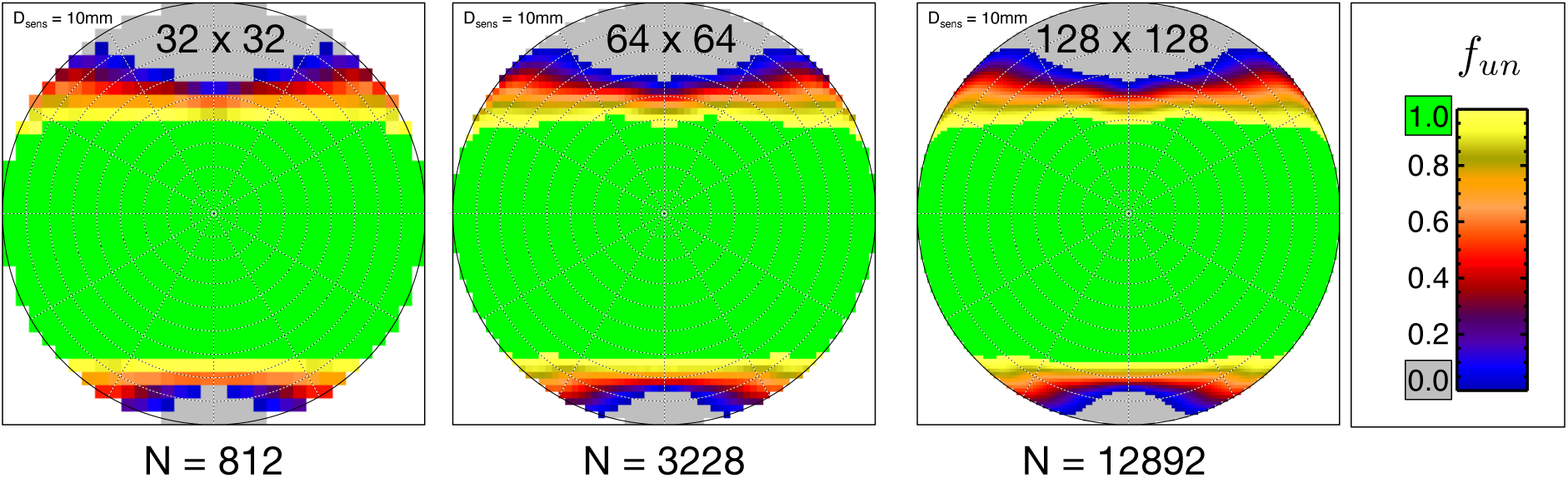
Example occlusion patterns for occlusion virtual goniophotometer (OVG) grid resolutions of 32×32, 64×64 and 128×128. *N* gives the number of light sources distributed across the OVG hemisphere.

**Figure 10:**
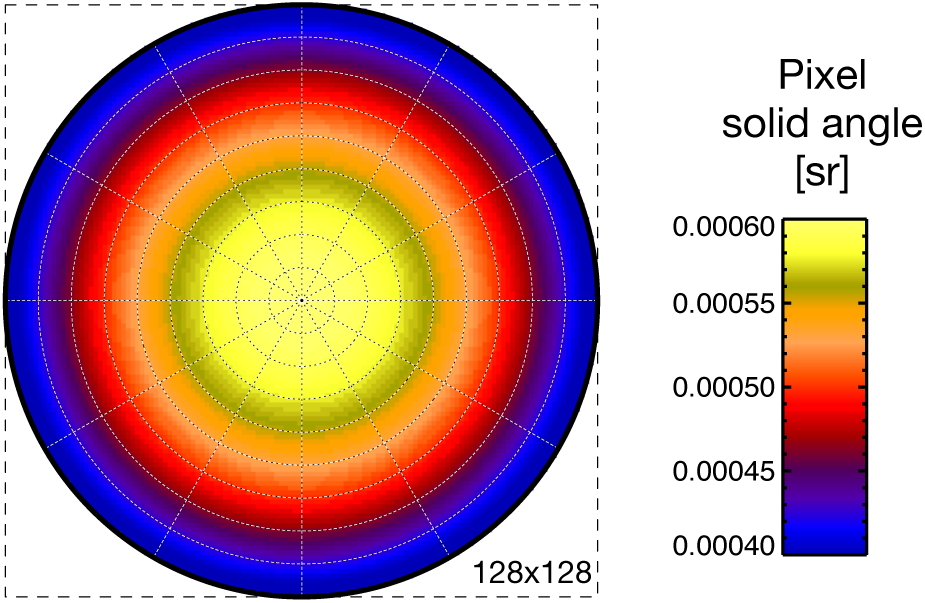
Pixel solid angle for the 128×128 for the occlusion virtual goniophotometer (OVG) hemisphere.

### 3.3 Measuring sensor disc response profiles

The effect of partial shading of a sensor disc will depend on the spatial responsivity of the sensor disc, i.e. how light at a certain position on the disc impacts the sensor’s reading. We refer to this quantity as the sensor (disc) response function. For the devices typically used by lighting researchers, it appears to be an entirely unknown quantity: we could find no information on measured sensor response functions, and the device manufacturers, when contacted, were unable to provide any data. We therefore devised our own approach to characterize it.

A laboratory experiment was conducted to determine the sensor response profiles of five commercially available light meters: a Konica Minolta CL500A, a Hagner E4-X, an Eltek LS50, a Konica Minolta T-10A, and a Testo 540. The latter two devices had a slightly curved sensor disc. The diameters of the sensor discs range from 8.4 mm to 26.5 mm. All devices had been previously calibrated for linear response, though only relative sensitivities were considered here, so absolute calibration is not relevant.

Specific areas of each sensor disc were illuminated sequentially by a narrow laser beam to determine the relative sensitivity across the disc. To accurately point the laser at a specific area of the sensor, it was mounted on a rail. The position of the laser on the rail was electronically controlled and could be moved in increments of 1 mm. Perpendicular to the rail, the light meter was clamped (Figure 11a). The clamp could be rotated by 90 degrees, thereby changing the light meter’s orientation and allowing the laser to be aimed along two axial directions across the sensor disc (Figure 11b). The number of measurements depended on the diameter of the sensor disc. For each laser position on the sensor disc, the (photopic) illuminance value measured by the light meter was recorded. All measurements were conducted in a completely dark room, except for illumination from the laser.

**Figure 11:**
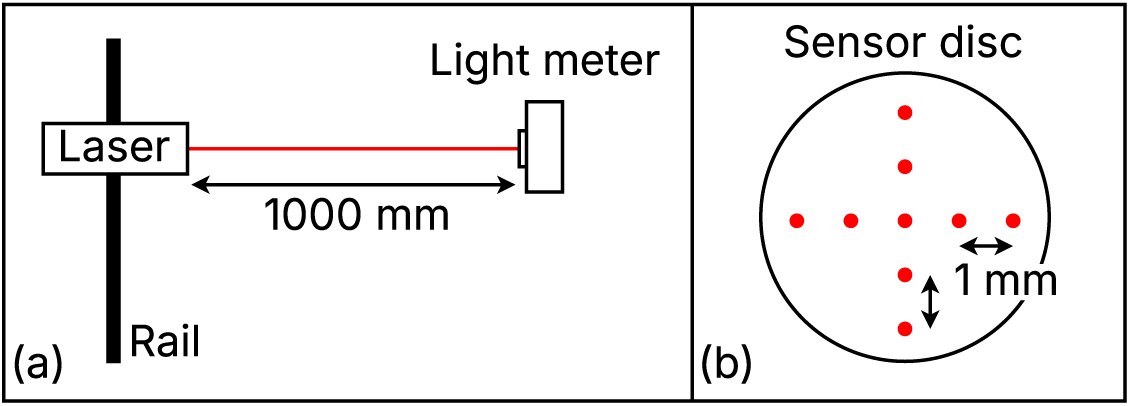
(a) Laboratory measurement setup to determine the sensitivity across a light meter’s sensor disc. The laser was aimed perpendicular to the sensor disc. (b) Positions where the laser beam was aimed at the sensor disc. The number of positions depends on the sensor disc’s size.

Two different lasers were used, a red laser (Laserfuchs LFD650-0.4-NT) with a wavelength of 650 nm and a green laser (TRU COMPONENTS LM05GND) with a wavelength of 532 nm. Both lasers were positioned such that the beam diameter at the sensor disc was approximately 2 mm. The red laser had an output power of 0.4 mW and provided illumination within the instrument range of the light meters. The green laser had a considerably higher output of 5.0 mW and was therefore dimmed to provide illumination within the instrument range of the light meters. The use of two different lasers was motivated by the very narrow spectral power distribution (SPD) of laser light. Although light meters are designed to measure radiation within the visible spectrum, with spectral sensitivity based on the V(*λ*) curve, and both lasers emit within this range, their SPDs are not representative of real-world daylighting or electric lighting SPDs. To account for potential wavelength-dependent effects resulting from using a light source with a narrow SPD, for example, differences in light scattering by the top (diffusing) layer of the sensor disc, measurements were therefore conducted using two lasers.

## 4 Results

The results are structured as follows. First, we present the measured sensor response profiles. Next, we describe how these are integrated into the OVG simulations. Finally, we present the results of the OVG simulations.

### 4.1 Measured sensor response profiles

The sensor response profiles for each light meter are shown in Figure 12. The illuminance measurements were normalized relative to the measurement when the laser beam was aimed at the center of the sensor disc. All sensor diameters were normalized from −1 to +1, with the center of the disc being at 0. The results from both light meter orientations, i.e. the measurements along two perpendicular axes over the sensor disc, were averaged. Since the results indicate a radial response over the sensor discs, the resulting values were averaged based on the distance from the disc’s center. No wavelength-dependent effects were observed between the results for each laser. Thus, based on these axially and mirror averaged measurements of both lasers, (radial) sensor response functions were determined for each light meter. The type of function was chosen based on the overall shape of the distribution of the measurement points, approximating a polynomial or Gaussian fit. All resulting radial sensitivity functions (Table 2) have an R^2^ value larger than 0.99.

**Figure 12:**
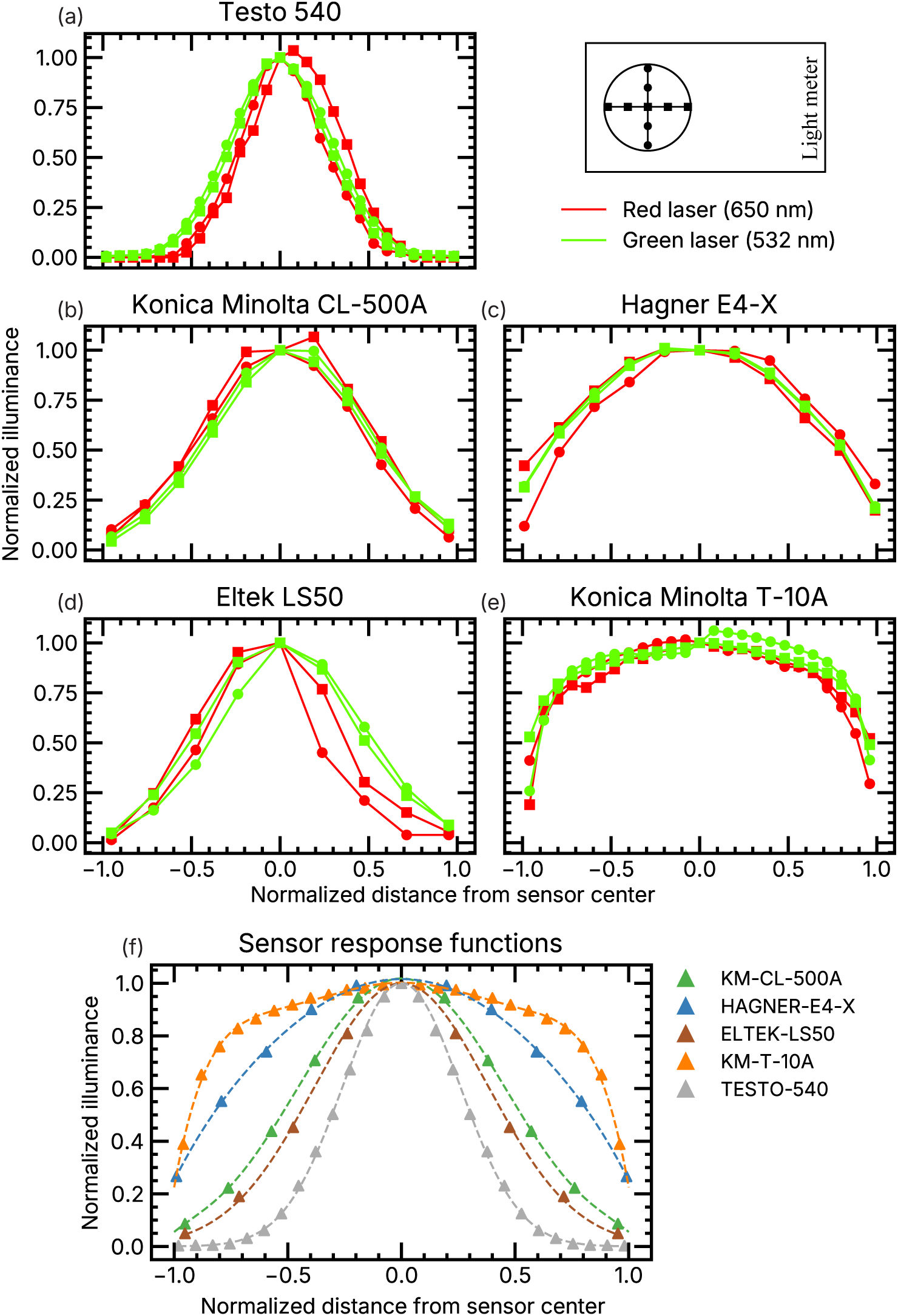
Measured line profiles of sensor disc response (a) to (e), and summary plot showing all five fitted sensor response functions (f).

**Table 2:** List of sensor response functions. As with Figure 12, normalized to sensor disc radius = 1.

| Device / Label | Normalized sensor response function |
| --- | --- |
| KM-CL-500A | $S = -0.88r^6 + 2.37r^4 - 2.45r^2 + 1.02$ |
| HAGNER-E4-X | $S = -0.14r^6 + 0.18r^4 - 0.80r^2 + 1.02$ |
| ELTEK-LS50 | $S = e^{(-3.42r^2)}$ |
| KM-T-10A | $S = -1.48r^6 + 1.31r^4 - 0.60r^2 + 1.00$ |
| TESTO-540 | $S = e^{(-7.45r^2)}$ |
| FLAT | $S = 1$ |
| TOP-HAT-1MM | $S = 1$ (modeled as FLAT 1 mm sensor) |

### 4.2 Application of measured sensor profiles to OVG simulations

Besides the five measurement-based sensor response functions (Figure 12), two theoretical functions were also evaluated (Table 2). FLAT, which assumes constant sensitivity across the disc, and TOP-HAT-1MM, which assumes a flat sensitivity across a 1 mm diameter region in the center of either the 7 mm or 10 mm discs, and zero elsewhere across the disc. This latter function was introduced when we learned additional detail regarding the sensor configuration behind the diffusing disc of the LIDO device: “The dimensions of the sensor’s opening behind the diffusor and the IR filter is approx. 1 mm” (K. Broszio, personal communication, 9^th^ March, 2026). The TOP-HAT-1MM response therefore is a simplification of the LIDO which assumes zero contribution to response outside of the 1 mm diameter center of the disc, and a flat response inside.

The vector **S** contains the list of the 656 sensor response values *S_i_* at each of the points calculated using any one of the response functions listed in Table 2. Independent of the sensor response function used, when the disc is evenly illuminated, the simulated ‘measurements’ from all functions should all agree. Thus, the sensor response vector needs to be normalized to **S_nor_** which is:

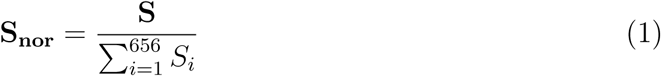

Although results were generated for all four occluders (CJ, PJ, CL, PL, Table 1) using all seven sensor response functions (Table 2), in the main body of the paper we present results for the three functions that span the range in performance outcomes; the remainder are available in Appendix A.2. The three are FLAT, TESTO-540, and TOP-HAT-1MM. The reason for this selection is that the differences between the various response functions shown in Figure 12 are smaller than they might appear once the increase in sensor area with radius is taken into account. For example, the outer tenth of the normalized radius (0.9 to 1) accounts for approximately 19% of the disc area, versus about 1% for the inner tenth (0 to 0.1). This is demonstrated by plotting each function as a cumulative effect with increasing radius, calculated using a finite element method that divides the normalized (0 to 1) radius disc into 1,000 annuli, Figure 13. It can then be seen that the FLAT and TOP-HAT-1MM functions represent the two extremes of sensor response, with TESTO-540 closest to a ‘middling’ response between them.

**Figure 13:**
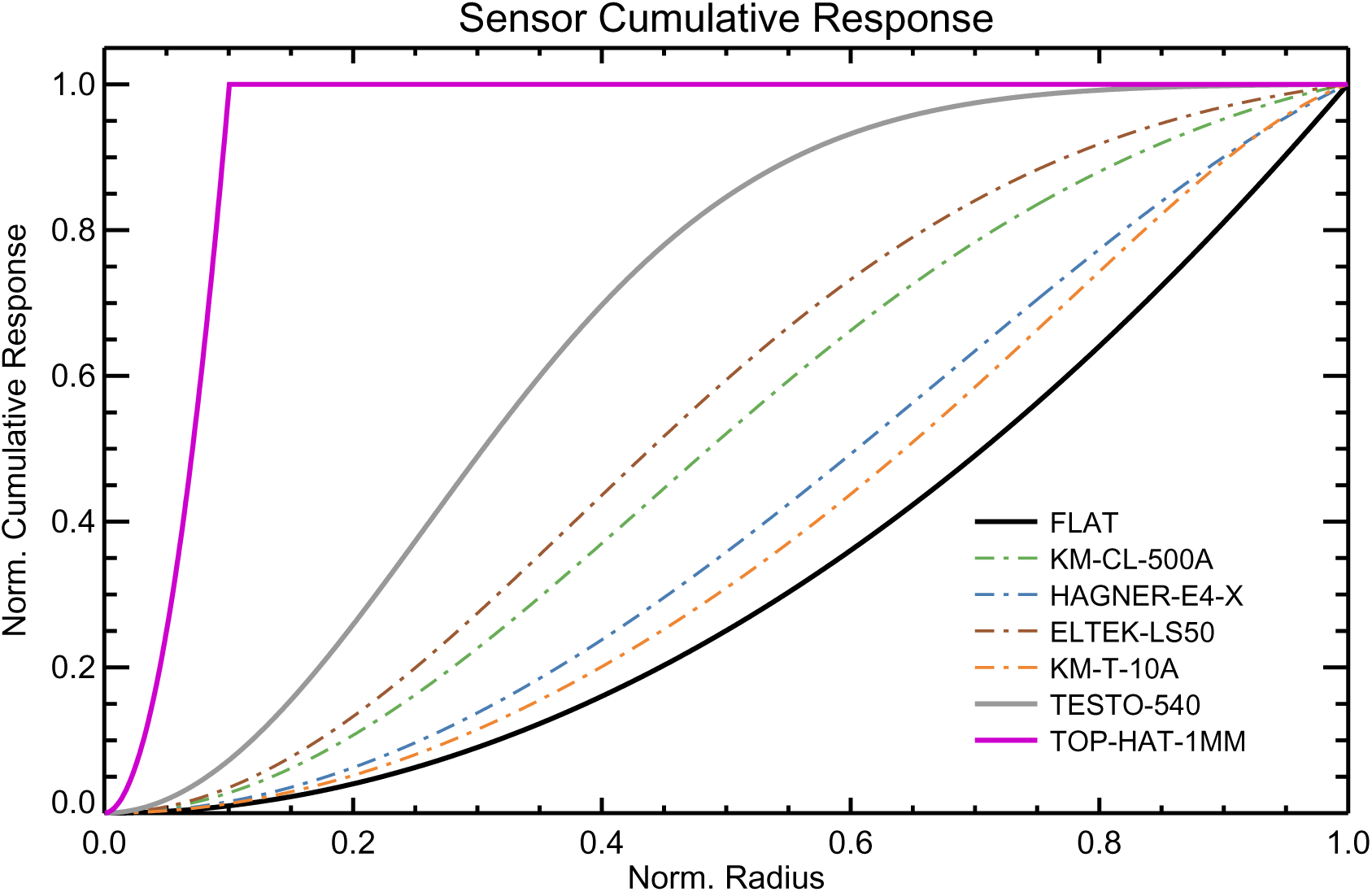
Cumulative sensor response with increasing (normalized) radius for the functions given in Table 2. The TOP-HAT-1MM curve is with respect to a 10 mm diameter sensor disc (the other curves are invariant to sensor disc diameter).

### 4.3 OVG results for JETI and LIDO devices

As the renderings of different viewpoints on the sensor discs of the JETI and LIDO (Figures 3 and 5) indicate, deviation from ideal CIE occluder behavior will have two effects on measurements depending on the direction from which the light originates:

1. Partial shading of the sensor from light originating within the CIE-defined FOV will result in a measurement that will be below the correct value, i.e. an underestimation.
2. Any illumination of the sensor (partial or full) from light originating outside of the CIE-defined FOV will produce an incorrect, greater than zero measurement, i.e. an overestimation.

To aid the understanding of the significance of the findings from the OVG, we present the results in terms of the absolute difference in irradiance between the performance of an ideal point sensor – one that follows the CIE definition exactly – and that of the JETI and LIDO devices fitted with the circumference-based (CJ, CL) and point-based (PJ, PL) occluders. Additionally, the outputs of each of the 12,892 point light sources in the OVG were normalized to deliver an irradiance of 100 Wm^-2^ on the (unshaded) plane of the sensor disc. Normalizing to a value of 100 (as the absolute units are not relevant in this context) allows for the *inference* of percentage errors in the CIE-defined FOV (where all values should equal 100). ‘Ideal’ performance of a sensor-occluder combination in the OVG is shown in Figure 14a, where the sensor is a single point, and the occluder is the point-based model. Values equal to 100 are shown in green, and those for zero are shaded gray. In Figure 14b, an example of predicted device performance of the CL occluder with an arbitrary measured profile (HAGNER-E4-X) applied as the sensor response function is shown. Values between zero and 100 are shaded according to the accompanying color scale. For approximately half of the FOV, the shading is green (100) or yellow (between 90 and *<*100). In other words, a region where underestimation is never greater than 10%. Orange shading (~70) within the FOV indicates an underestimation of ~30%. Outside of the FOV, shading reveals a non-zero value which, ideally, should be zero. In the (ideally) occluded regions, the interpretation is slightly different from that used in the FOV. For example, the orange shading just showing in the upper occluded region shows a value of ~70 (where it ideally should be zero). This indicates that, for light incident from those directions, the measurement would be ~70% of the (unshaded) value that would have resulted from those (arbitrary) light sources.

**Figure 14:**
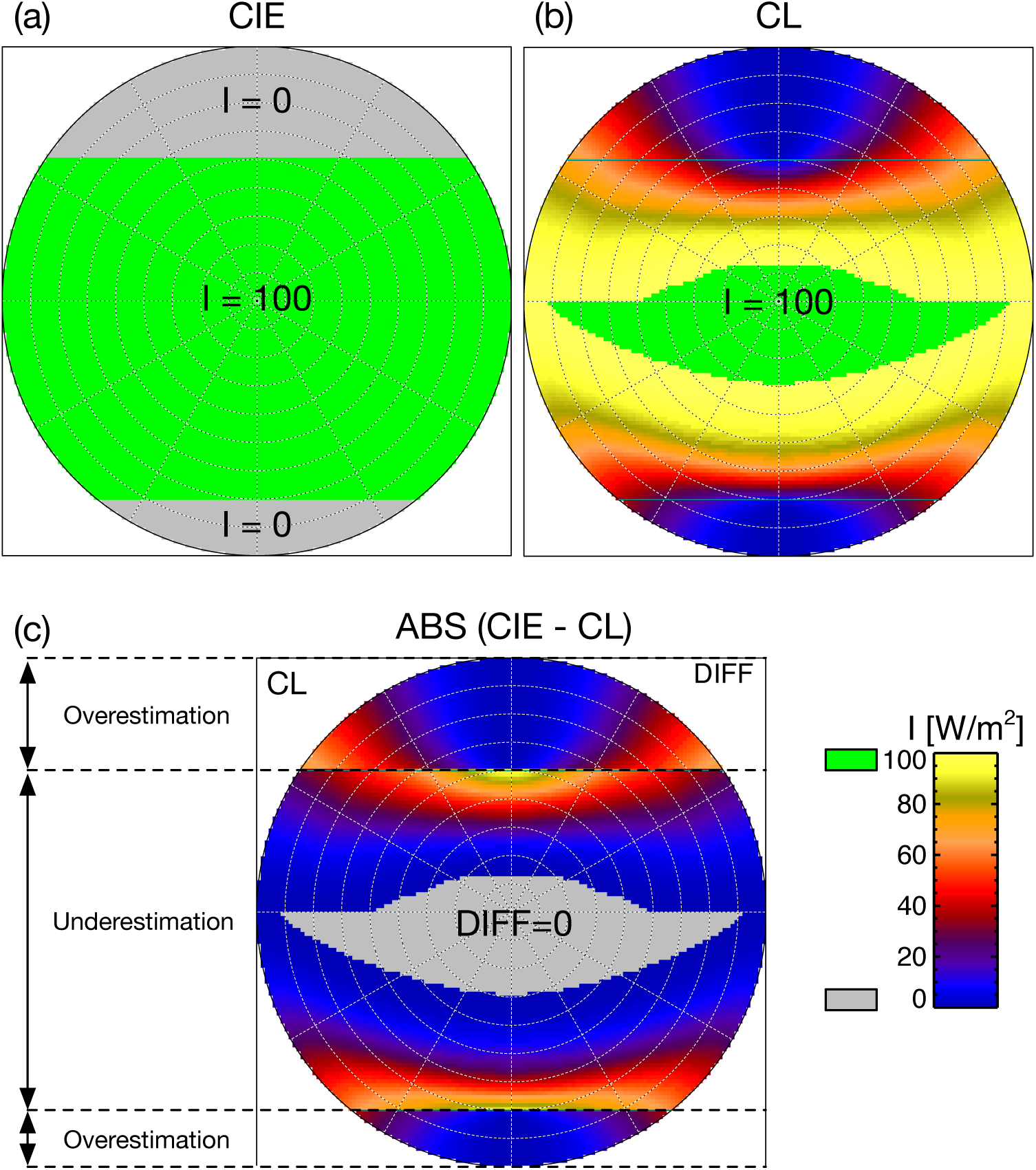
(a) occlusion virtual goniophotometer (OVG) results for a point sensor (i.e. the sensor disc is infinitely small) with a point-based occluder model, i.e. ‘ideal’ results based on the CIE S 026:2018 definition. (b) Results for the CL occluder with an example applied sensor response function (HAGNER-E4-X). (c) Absolute difference between (a) and (b).

The interpretation of Figure 14b with regard to the use of the false-color scale ‘flips’ depending on the (CIE-defined) region: the FOV region, or the two occluded regions. That necessity can be avoided by presenting the results of sub-figure (b) in terms of their absolute difference with the (ideal) results shown in sub-figure (a). This ‘difference’ plot is shown in Figure 14c. The advantage of sub-figure (c) is that a single reading rule applies everywhere: gray marks agreement with the ideal (zero difference), and any color marks a measurement error, regardless of region. The two occluded regions in sub-figure (c) are identical to those in sub-figure (b) because those regions are zero in the ideal performance plot, sub-figure (a). For the FOV region, gray therefore now marks zero difference rather than full occlusion, so relative to sub-figure (b) the scale is effectively inverted there. In this region, the irradiance value now relates directly to the percentage by which the measurement using the CL occluder underestimates the ideal, CIE-defined measurement. For example, the blue shade (in the FOV region) indicates that, for light incident from those directions, the measurement error will be about −10%, i.e. an underestimation. Red and orange shades indicate a measurement error of about −50% and −70%, respectively. For the two occluded regions, any shading that is not gray now indicates an overestimation (i.e. *>* 0) measurement for light originating from those directions (as described above).

The OVG results for the JETI and LIDO occluders are shown in Figures 15 and 16, respectively. In each figure, the first row shows the results using the FLAT sensor response function, sub-figures (a) and (b); the second row the TESTO-540 function, sub-figures (c) and (d); and, the third row the TOP-HAT-1mm function, sub-figures (e) and (f).

**Figure 15:**
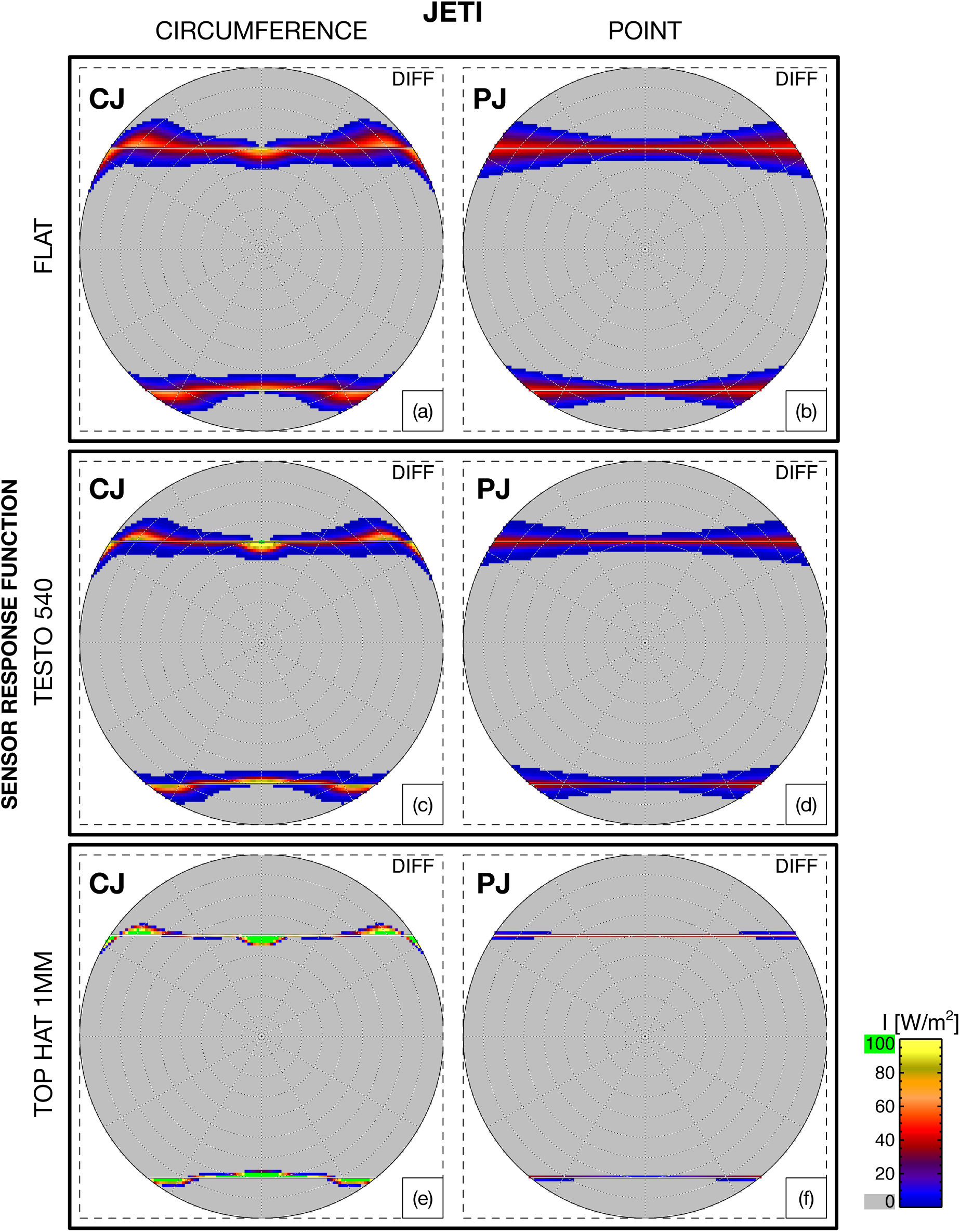
Occlusion virtual goniophotometer (OVG) absolute differences between the CIE S 026:2018 definition and the JETI occluders. Panels pair an occluder type – circumference-based (CJ) or point-based (PJ) – with a sensor response function: (a) CJ, FLAT; (b) PJ, FLAT; (c) CJ, TESTO-540; (d) PJ, TESTO-540; (e) CJ, TOP-HAT-1MM; (f) PJ, TOP-HAT-1MM.

**Figure 16:**
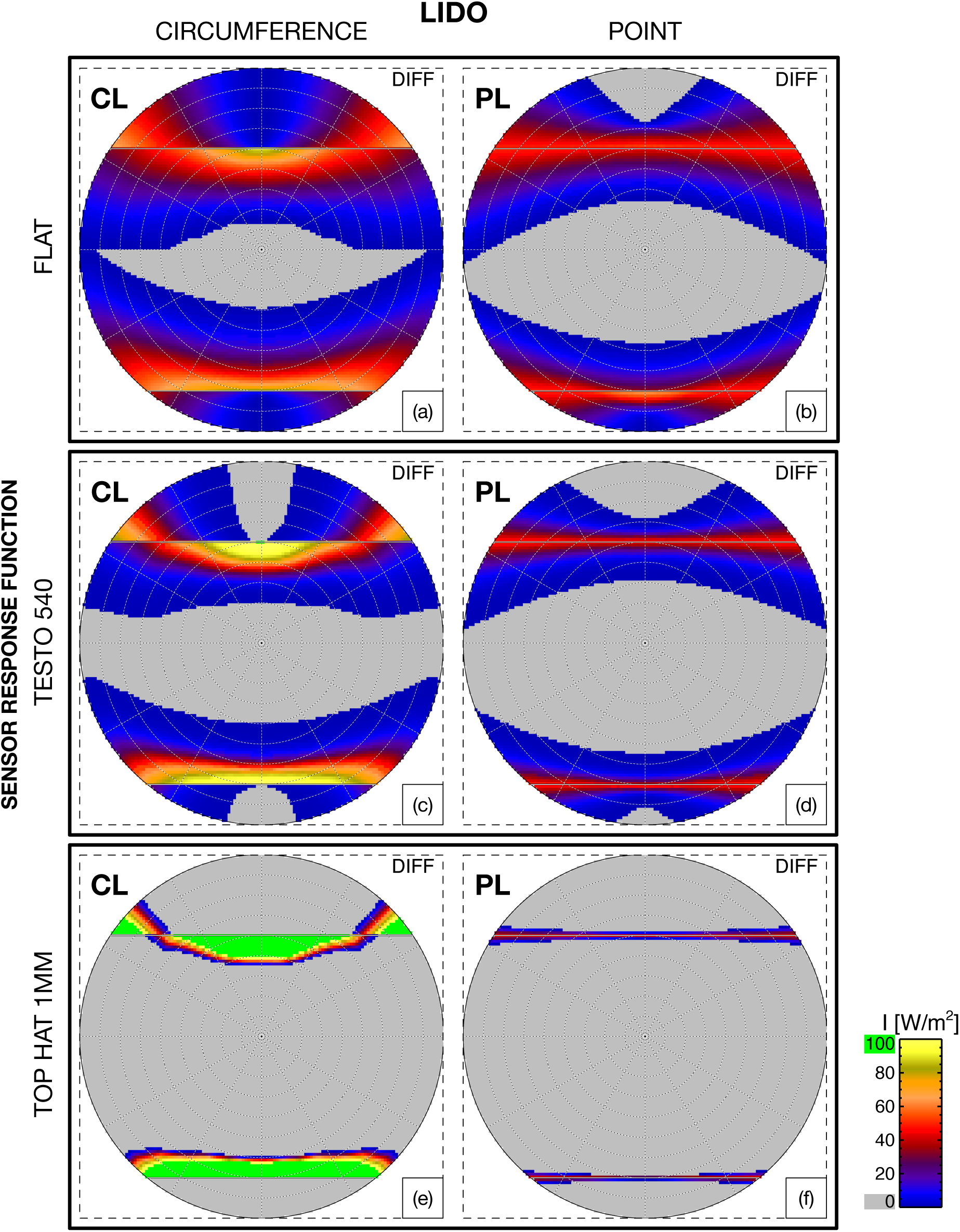
Occlusion virtual goniophotometer (OVG) absolute differences between the CIE S 026:2018 definition and the LIDO occluders. Panels pair an occluder type – circumference-based (CL) or point-based (PL) – with a sensor response function: (a) CL, FLAT; (b) PL, FLAT; (c) CL, TESTO-540; (d) PL, TESTO-540; (e) CL, TOP-HAT-1MM; (f) PL, TOP-HAT-1MM.

## 5 Discussion

The results presented in Figures 15 and 16 show that, irrespective of the applied sensor response functions, the point-based occluder model always performs equivalently or better than the circumference-based occluder model. Performance between the two occluder types is largely equivalent when the ratio of the occluder radius to the *effective* radius of the bulk of the disc sensitivity is around five or greater (the aforementioned ‘five times rule’). When this holds, the potential for partial shading across the sensor disc is greatly reduced. Furthermore, the shapes of the circumference and point occluders will ‘appear’ very similar from any point on the disc. As this is the case for the JETI occluders (ratio of ~5.14, Table 1), the circumference-based (CJ) and point-based (PJ) occluder models perform similarly across all pairs (Figure 15). And, overall, the agreement with the ideal CIE definition is good over the majority of possible incident light directions. However, it is clear that where divergence from ideal performance does occur (i.e. any non-gray shading), the circumference-based occluder model always performs less well than the simpler point-based occluder model. This is evident at the two boundaries between (ideal) FOV and (ideal) occlusion, e.g. yellow and even some green shading indicating 100% divergence from ideal.

The OVG results for the LIDO occluders show larger deviations, Figure 16. For the LIDO, the occluder radius is only about ~1.6× the sensor radius (Table 1). As expected for the FLAT sensor model, the effect of partial illumination of the sensor (i.e. non-gray shading) is evident for the majority of possible incident light directions – less than half of the hemisphere of possible directions is shaded gray, sub-figures (a) and (b). Nevertheless, the point-based occluder model (PL), sub-figure (b), shows markedly less divergence from ideal than the circumference-based occluder model (CL), sub-figure (a). Applying the TESTO-540 sensor response function, the divergence from ideal FOV for the CL occluder is increased at the FOV boundaries, i.e. regions of orange/yellow shading in sub-figure (c). Whereas for the PL occluder the divergences from ideal are lower in both extent and magnitude, sub-figure (d). Lastly, the results using the TOP HAT 1mm function are shown in sub-figures (e) and (f). For the CL occluder the divergences reach 100 (green shading) at both FOV boundaries (e). This means that, for the incident light directions shaded green inside the (ideal) FOV, the sensor response will be zero. For both the upper and lower regions of the FOV the green shading achieves an angular extent of ~10°. Thus, at the vertical meridian, the upper and lower cut-off angles will be around 40*°* and 60*°*, respectively, rather than the 50*°* and 70*°*angles in the CIE definition. In contrast, the PL occluder (f) exhibits near ideal performance with only narrow regions of low divergence (i.e. mostly blue shading) along the FOV boundaries.

How the divergences from ideal occluding performance manifest as measurement errors in real settings will depend strongly on the nature of the luminous environment, in particular the angular size and distribution of light sources in the FOV. The authors have carried out a number of simulations to gauge the effect of the different occluder-sensor combinations. For example, traversing different-sized light sources across 180*°* arcs in 5*°* increments centered on the occluder. However, the outcomes of such tests are heavily dependent on how the test is configured, and so are likely to be potentially misleading if attempts are made to generalize the findings in terms of summary performance metrics. Additionally, such tests reveal little that cannot be inferred from the OVG difference plots.

To illustrate how differences in FOV occlusion between the PL and CL occluders can lead to measurement discrepancies, a simulated indoor environment is shown in Figure 17. The rendered images left and right are identical. The rendering was generated using a 180*°* angular fisheye projection. Thus, the individual OVG difference plots shown in Figures 15 and 16 can be superposed directly on to the renderings. For this, the gray areas of zero difference in the OVG plots were made fully transparent, and the remaining shades partially transparent. The OVG difference plots for the CL and PL occluders, calculated using the TOP-HAT-1mm response function (bottom row in Figure 16), were then superposed on the same rendering, left and right, respectively. The composed images therefore show the dosimeters’ ‘view’ of the scene with the directional difference in performance from the ideal shown by the (semi-transparent) shading of the OVG plot. Depending on their light output distribution, ceiling luminaires directly in the FOV of the light meter can be significant contributors to the measured illumination at that point. Three ceiling luminaires labeled P, Q and R have been highlighted in both images. Luminaires P and Q are fully inside the CIE-defined FOV, and luminaire R is fully outside of the FOV. For the CL occluder, the P and Q luminaires (which should be inside the FOV) are showing as significantly occluded: P is showing as almost fully occluded (mostly green shade) and the extent of Q varies from fully occluded (green) to just unoccluded at the edge. Whereas luminaire R, which is fully outside of the FOV, varies from fully visible (green) to just fully obstructed at the top edge. From this viewpoint, the illumination contribution of P will be almost zero, whilst from Q it will be significantly reduced. Luminaire R, where the illuminance contribution should be zero, will contribute about half of what it would if the CL occluder were to be removed. In contrast, with the PL occluder the luminaires P and Q are (correctly) fully unobstructed. Luminaire R (outside of the FOV) is nearly fully obstructed – there is just a tiny portion of R shaded blue, i.e. only ~10% of that small part of the luminaire would contribute a small amount of illuminance to the total from the rest of scene in the FOV.

**Figure 17:**
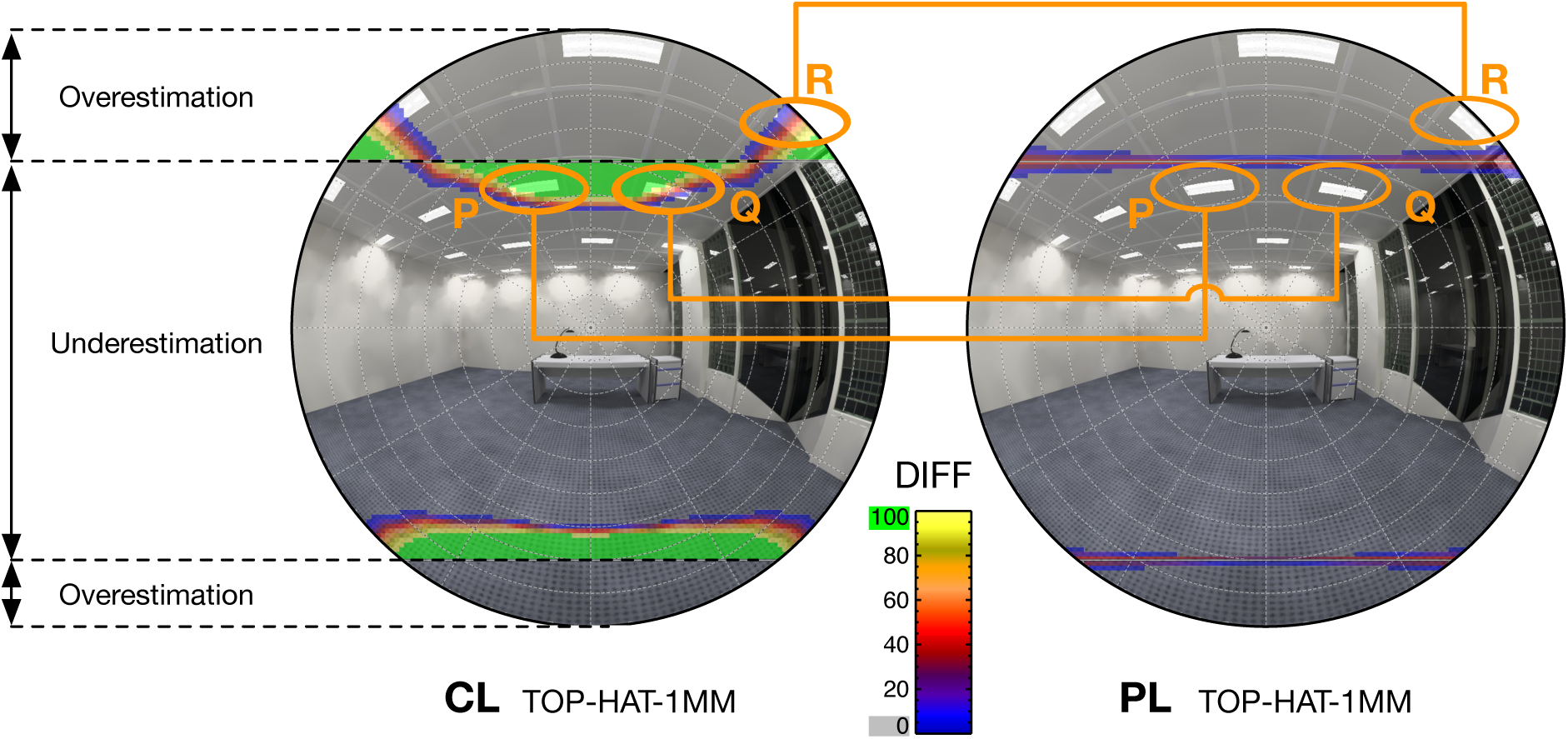
Hemispherical fields of views (FOVs) of a LIDO equipped with the point-based (PL) and circumference-based (CL) occluders in an example indoor environment. Differences in occlusion performance between each and the CIE ideal are shown in false color. Labels P, Q, and R refer to luminaires in the scene.

The viewpoint used in Figure 17 clearly illustrates the superior performance of the point-based LIDO occluder (PL) compared to the original circumference-based (CL) occluder. However, with only a small change in position, one could obtain a view where none of the false-color in the CL difference plot overlaps with any ceiling luminaire. The light arriving directly from the luminaires would then be correctly accounted for, masking the better performance of the point-based occluder. Consequently, we propose it is more reliable to compare *intrinsic* performance based on the OVG results and with respect to the CIE definition, rather than measurements (real or simulated) in any actual space. This is simply because the OVG gives information on *all* possible incident light directions independently of each other, whereas the outcome from measurements taken in any particular environment will be highly specific to that environment. There is a direct parallel with, say, comparing two photometers: all other things being equal, the photometer with the better cosine response curve is the one to be preferred.

To our knowledge, no prior studies have used goniophotometry to characterize the effects of partial occlusion of light meters, or empirically determined sensor response functions. The results presented here therefore cannot be compared directly with prior work.

### 5.1 Limitations

The OVG approach has, in our view, no intrinsic limitations that might restrict reasonable expectations of accuracy. Inherent precision is directly related to the OVG grid dimensions – this can be increased as required. The 128×128 grid gave 12,892 light sources distributed across the hemisphere. Grids of 256×256 and 512×512 would produce 51,468 and 205,892 light sources, respectively. With each quadrupling of light sources there is an approximate halving of the angular separation between the sources. The 3D occluder models used in the simulations are identical to those used by the 3D printer to create the physical occluders. Though, one factor outside of our control is the degree to which a 3D-printed occluder exactly matches the 3D geometry ‘sent’ to the printer.

The experimental setup for characterizing sensor response profiles (Figure 11) requires a high level of precision, as the laser must be pointed at the center of the sensor disc. Because the light meter cases had different dimensions, clamping them required manually locating the center of each sensor disc. As a result, a slight shift in the center position is visible in Figure 12 for some measurements. However, such shifts are not expected to have affected the resulting sensor response functions, as measurements from both lasers were axially and mirror averaged before fitting the functions.

The sensor response functions were characterized for five commercially available light meters, which do not include the JETI or LIDO devices. The actual response functions of these two devices thus remain unknown. Since the OVG results include, in addition to these five response functions, two functions representing boundary cases (FLAT and TOP-HAT-1MM), the results span a plausible range of sensor responses that is expected to include the actual JETI and LIDO functions. However, the absolute divergence for any particular device would require characterizing that device’s response. Additionally, two of the considered devices (Konica Minolta T-10A and Testo 540) have slightly curved sensor discs, whereas the OVG assumes a flat disc. Hence, actual performance may diverge from the OVG results for such devices.

Lastly, measurements with a FOV occluder only provide an approximation of retinal light exposure since light is further modified as it passes through the ocular media, i.e. cornea, aqueous humor, lens, and vitreous humor. Incorporating such aspects in measurements is not straightforward, as they are influenced by factors such as the pupillary size, lens characteristics, and gaze direction of the pupils, which are prone to differ due to inter-individual differences and the surrounding illumination condition (van Derlofske et al. 2000; Watson and Yellott 2012; Spitschan et al. 2022; He et al. 2024). Additionally, studies have shown that where the light is originating from within the human FOV, i.e. the incident angle at the cornea, may affect the magnitude of (nighttime) NIF-related responses (Visser et al. 1999; Lasko et al. 1999; Smith et al. 2002; Glickman et al. 2003; Rüger et al. 2005; Rea et al. 2021). Thus, to consider such effects, it may be necessary to perform spatially resolved measurements (Broszio et al. 2018; Knoop et al. 2019).

### 5.2 Recommendations and code availability

When occluding part of a light meter’s FOV to represent the human FOV according to the CIE S 026:2018 definition, we recommend an occluder model based solely on the occluder radius, i.e. a point-based occluder. This is both the simplest approach and performs at least as well as the considered circumference-based models. When significant differences were found, the point-based model always performed better than the circumference-based models. In a point-based model, the radius of the sensor disc only matters for determining the occluder’s radius. A ratio of ~5 is generally adequate, meaning the occluder radius should be at least five times the sensor disc radius – or, if the sensor response function is known, at least five times the radius of the region with effective sensitivity. When the occluder radius can be made large enough to satisfy this ratio (relative to the sensor disc radius), the actual sensor response function has a relatively minor influence on the measurements. If this is not possible due to design constraints, it may be worthwhile to characterize the sensor response function, or to approximate it using one of the sensor response functions in Figure 13. It *may* then be possible to estimate the divergence from CIE-defined FOV performance using the results presented here. Otherwise, it may be necessary to evaluate the particular occluder-sensor configuration using an OVG similar to the one described here.

The script used to model the point-based occluders is openly available (de Vries and Mardaljevic 2026) and supports two use cases. For lighting simulations, it generates a *Radiance* geometry definition that can be used directly, as demonstrated in previous studies (de Vries et al. 2026b). The occluder geometry consists only of two walls made of facets (see Figure 4a for an example). Because light sensors are typically modeled as point sources in such simulations, the occluder model is exact, and its absolute size is irrelevant. For use with actual light meters, the script generates a watertight mesh suitable for 3D printing. Since absolute size matters here, the user can set the occluder radius to satisfy the 5× ratio for a given sensor disc. Additional parameters control the thickness of the occluder walls and whether a ring connects the upper and lower occluder. Examples of 3D-printed occluders using this script are available in Appendix A.3.

## 6 Conclusion

The study described in this article is, to our knowledge, the first to rigorously evaluate the performance of occluders designed to represent the FOV definition for corneal light exposure specified by the CIE. The method developed is a simulation-based approach founded on goniophotometric principles. Our method assessed compliance with the CIE definition for all possible incident light directions to give a comprehensive appraisal of the intrinsic performance of the occluder. Two occluder types were evaluated: the circumference-based model originally proposed by Zauner et al. (2023), and the simpler point-based model, which the authors derived from the CIE definition analytically.

Our approach revealed that the sensor disc response function could be an additional significant factor influencing the measure of compliance with the CIE definition. This is possible when the occluder radius is not much greater than the sensor disc radius, as is likely for compact devices such as the LIDO. The response function of the light sensor discs appeared to be a previously undetermined quantity. This necessitated the development of an ad hoc experimental technique using lasers to measure the sensor response at increments across the diameter of the sensor disc. In addition to the measured sensor response functions, we added two limiting cases which we consider to encapsulate the range likely to be found in most devices.

The results showed that the most important factor determining compliance is the ratio of the occluder radius to that of the sensor disc. If that ratio is ~5 or greater (e.g. JETI device), both the circumference-based and point-based occluders show good compliance with the CIE definition. Additionally, any effect on compliance due to the disc sensor response function was found to be small across the full range of tested functions, i.e. from FLAT to TOP-HAT-1MM. However, although the divergence in performance between the circumference-based and point-based occluders was fairly small, the point-based occluder consistently showed higher compliance with the CIE definition.

For the LIDO device, the ratio of the occluder radius to the sensor disc radius was ~1.6, and the difference in performance between the circumference-based and point-based occluders was much greater than that observed for the JETI device. The point-based occluder delivered a higher degree of compliance with the CIE definition than that observed for the circumference-based occluder across the full range of sensor response functions tested. We therefore recommend occluders designed using the point-based model to represent the human FOV, irrespective of the ratio in occluder to disc radii. To support the use of such occluders in research and practice, we provide accompanying code that generates both *Radiance* geometry for lighting simulations and watertight meshes for 3D printing to be used as physical occluders on actual light meters.

## Appendix

### A.1 Occluded FOV angles for three positions on the sensor discs

The solid angles that comprise the FOVs shown in the parallax renderings (Figures 3 and 5) are given in Tables A1 and A2, for the JETI and LIDO occluders, respectively. For the JETI, the numerical differences in solid angles between respective CJ and PJ pairs is very small, probably below the threshold of practical significance. For the LIDO however, the differences are much larger. Consider the FOV solid angles from the −5.0 mm position against the correct value of 5.011 sr (PL from the center). For the CL occluder the FOV is 3.523 sr, whereas for the PL it is 4.508 sr – almost a difference of 1 sr. Note, such illustrative divergences in solid angles are only *suggestive* of performance differences.

**Table A1:** Subtended solid angles of field of view (FOV) and occluded view for CJ and PJ occluders from the three view positions shown in Figure 3. The values in bold correspond to the FOVs that exactly match the CIE S 026:2018 definition.

| Occluder Type | View point y shift [mm] | FOV solid angle [sr] | Upper occluder solid angle [sr] | Lower occluder solid angle [sr] |
| --- | --- | --- | --- | --- |
| CJ | 0.0 | 5.005 | 0.949 | 0.330 |
|  | +3.5 | 4.769 | 1.277 | 0.238 |
|  | -3.5 | 5.081 | 0.720 | 0.482 |
| PJ | **0.0** | **5.011** | **0.947** | **0.326** |
|  | +3.5 | 4.776 | 1.274 | 0.233 |
|  | -3.5 | 5.096 | 0.712 | 0.476 |

**Table A2:** Subtended solid angles of field of view (FOV) and occluded view for CL and PL occluders from the three view positions shown in Figure 5. The values in bold correspond to the FOVs that exactly match the CIE S 026:2018 definition.

| Occluder Type | View point y shift [mm] | FOV solid angle [sr] | Upper occluder solid angle [sr] | Lower occluder solid angle [sr] |
| --- | --- | --- | --- | --- |
| CL | 0.0 | 4.571 | 1.106 | 0.606 |
|  | +5.0 | 3.155 | 2.889 | 0.239 |
|  | -5.0 | 3.523 | 0.503 | 2.258 |
| PL | **0.0** | **5.011** | **0.947** | **0.326** |
|  | +5.0 | 3.685 | 2.451 | 0.148 |
|  | -5.0 | 4.508 | 0.412 | 1.364 |

### A.2 OVG results for remaining sensor response profiles

The OVG results for the JETI and LIDO occluder configurations with sensor response functions KM-CL-500A, HAGNER-E4-X, ELTEK-LS50 and KM-T-10A are shown in Figure A1 (JETI) and Figure A2 (LIDO).

**Figure A1:**
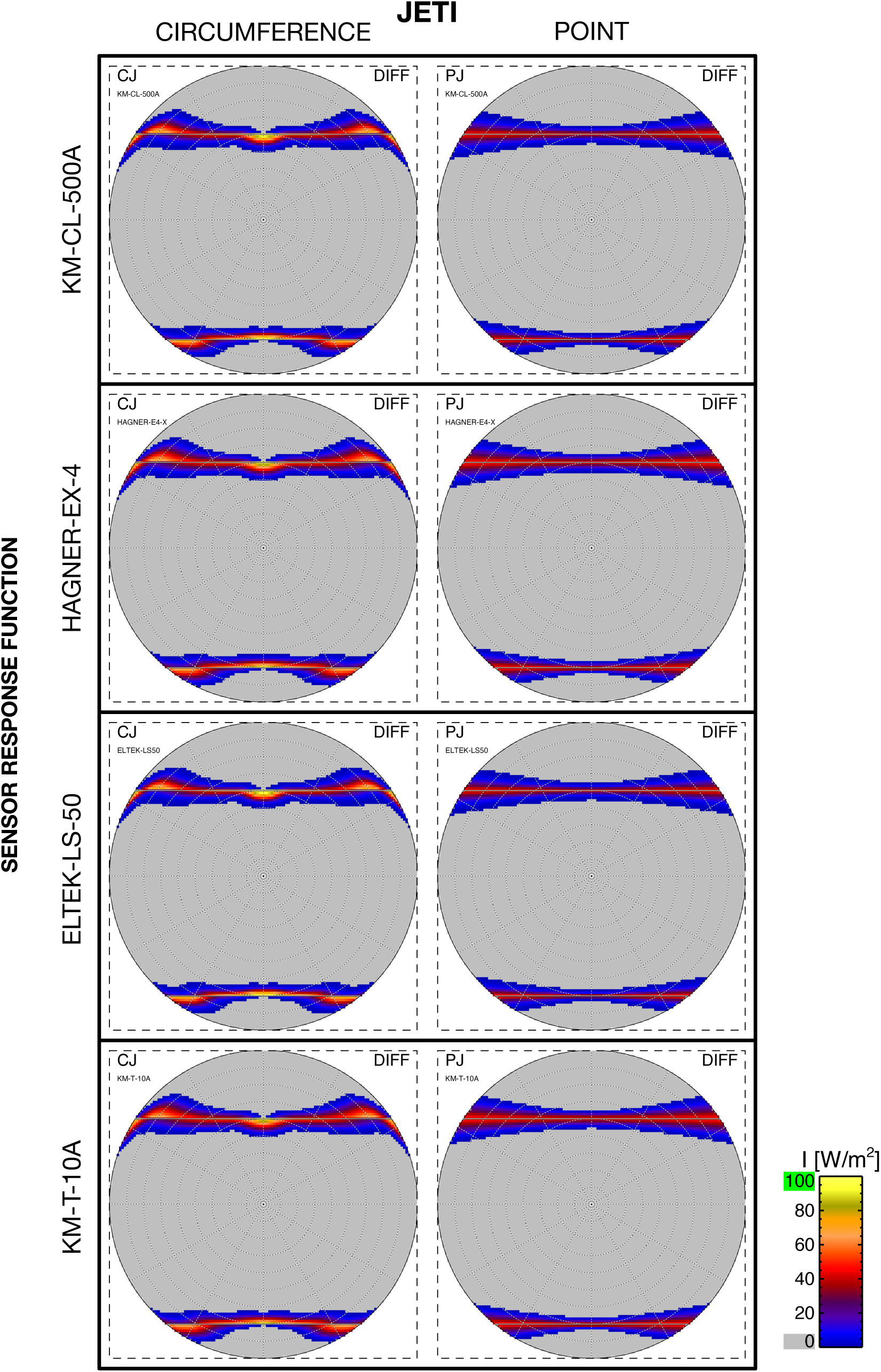
JETI circumference-based (CJ) or point-based (PJ) occluder difference plots

**Figure A2:**
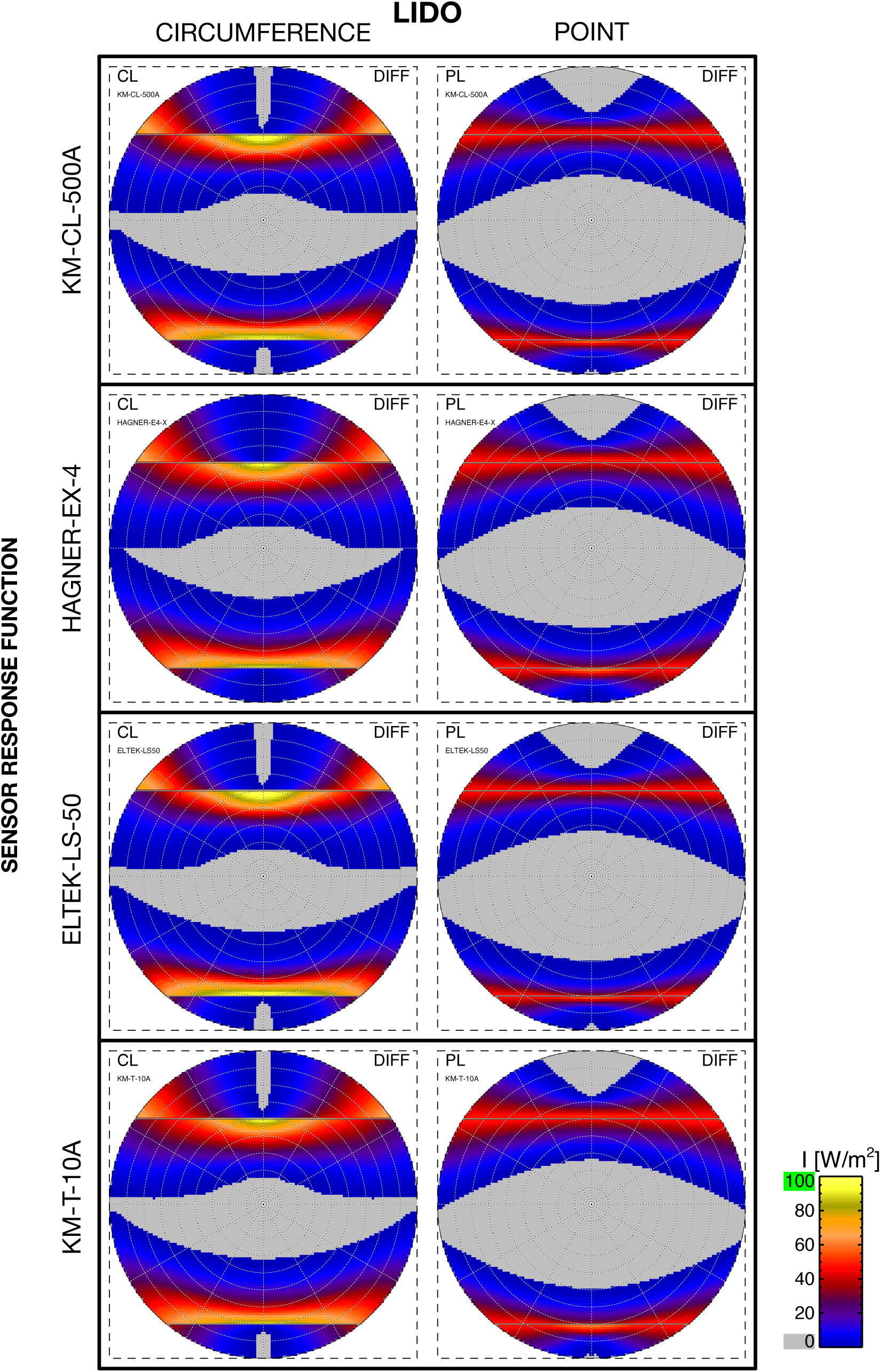
LIDO circumference-based (CL) or point-based (PL) occluder difference plots

### A.3 Examples of 3D-printed point-based occluders

3D-printed FOV occluders for the Konica Minolta CL500A and Hagner E4-X are shown in Figure A3. The occluders were generated using the point-based approach via an openly available script (de Vries and Mardaljevic 2026). For each occluder, the radius of the occluder equals five times the radius of the sensor disc.

**Figure A3:**
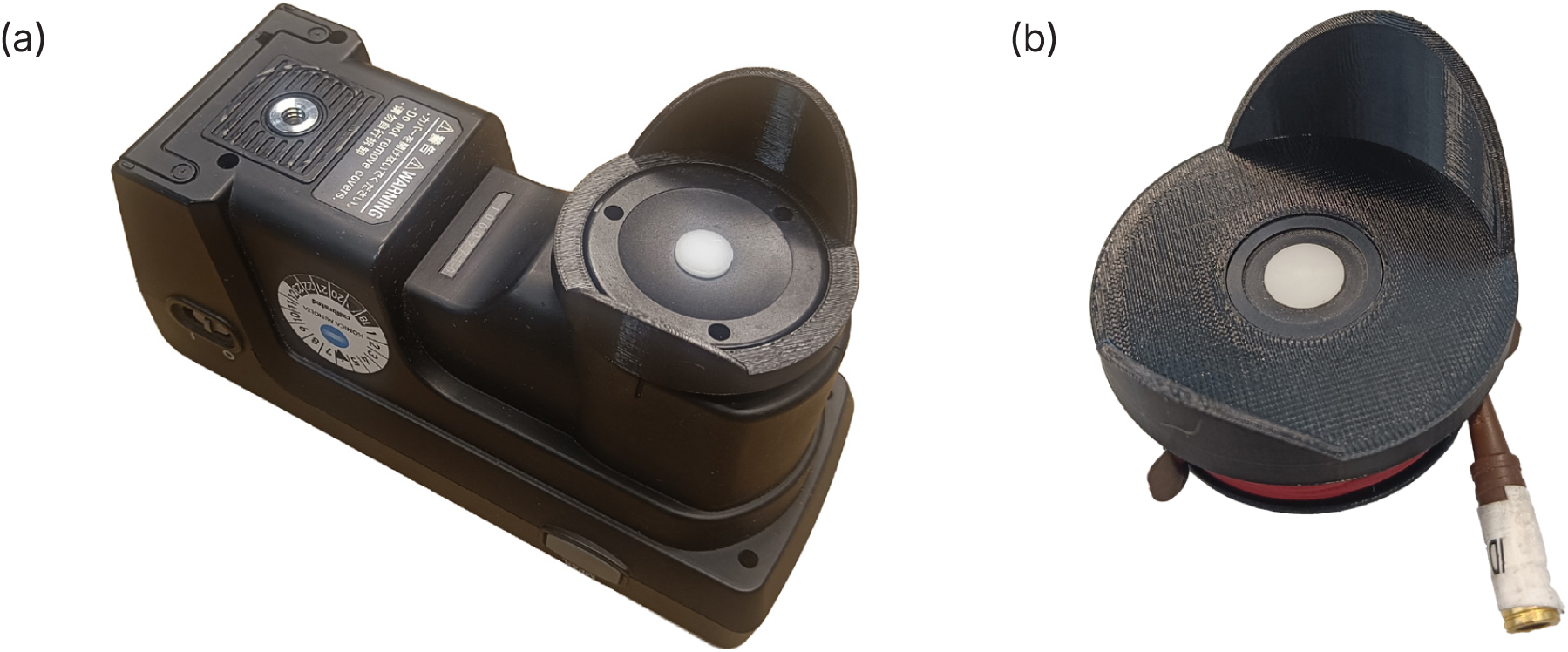
3D-printed (point-based) FOV occluder for the Konica Minolta CL500A (a) and Hagner E4-X (b).

## Acknowledgments

The authors wish to thank Johannes Zauner and Kai Broszio for supplying information on, and 3D occluder models of, the JETI and LIDO devices, respectively.

## Funding statement

No funding was received for this research.

## Disclosure statement

The authors report there are no competing interests to declare.

## Data availability statement

The datasets generated and analyzed during the current study are available from the corresponding author on reasonable request. The script used to generate the point-based occluders is available in an online repository (https://github.com/SietsedeVriesTUe/point-based-occluder) and as an archived release (de Vries and Mardaljevic 2026).

## Declaration of generative AI use

SWV used *Grammarly* and *Anthropic Claude Opus 4.8* to enhance the readability of the text. The authors reviewed the final manuscript and take full responsibility for its content.

## Author contributions

**JM:** Conceptualization, Formal analysis, Methodology, Software, Visualization, Writing – Original Draft, Writing – Review & Editing

**SWV:** Conceptualization, Formal analysis, Investigation, Methodology, Software, Visualization, Writing – Original Draft, Writing – Review & Editing

**JD:** Conceptualization, Supervision, Writing – Review & Editing

## Notes

https://doi.org/10.4121/a3b8c37a-de2d-48e4-9eee-ce456a2f98ce

### Competing Interest Statement

The authors have declared no competing interest.

## References

Alight A, Jakubiec J. 2026. An ipRGC-influenced/non-visual spectral occupant model (iNSOM) for lighting design, part 1: light simulation method. Lighting Res Technol. 58(1-2):6–22. 10.1177/14771535251368379.

Andersen M, Mardaljevic J, Lockley SW. 2012. A framework for predicting the non-visual effects of daylight – part I: photobiology-based model. Lighting Res Technol. 44(1):37–53. 10.1177/1477153511435961.

Apian-Bennewitz P, von der Hardt J. 1998. Enhancing and calibrating a goniophotometer. Sol Energy Mater Sol Cells. 54(1-4):309–322. 10.1016/S0927-0248(98)00082-8.

Appelfeld D, McNeil A, Svendsen S. 2012. An hourly based performance comparison of an integrated micro-structural perforated shading screen with standard shading systems. Energy Build. 50:166–176. 10.1016/j.enbuild.2012.03.038.

Bartell FO, Dereniak EL, Wolfe WL. 1981. The theory and measurement of bidirectional reflectance distribution function (Brdf) and bidirectional transmittance distribution function (BTDF). Radiation Scattering in Optical Systems. International Society for Optics and Photonics. SPIE. p 154 – 160. 10.1117/12.959611.

Bertenshaw DR. 2020. The standardisation of light and photometry – a historical review. Lighting Res Technol. 52(7):816–848. 10.1177/1477153520904755.

Blume C, Garbazza C, Spitschan M. 2019. Effects of light on human circadian rhythms, sleep and mood. Somnologie. 23(3):147–156. 10.1007/s11818-019-00215-x.

British Standards Institute. 2005. BS 667:2005 Illuminance meters – Requirements and test methods. British Standards Institution, London.

Broszio K, Knoop M, Niedling M, Völker S. 2018. Effective radiant flux for non-image forming effects: is the illuminance and the melanopic irradiance at the eye really the right measure? Light Eng. 26(2):68–74. 10.33383/2018-003.

Broszio K et al. 2025. Optimization of personal light dosimetry: consideration of the binocular field of view. In: Proceedings of the CIE Midterm Meeting 2025. Vienna (Austria): CIE. 10.25039/x051.2025.

Brown TM et al. 2022. Recommendations for daytime, evening, and nighttime indoor light exposure to best support physiology, sleep, and wakefulness in healthy adults. PLOS Biol. 20(3):e3001571. 10.1371/journal.pbio.3001571.

Danell M, Ámundadóttir ML, Rockcastle S. 2020. Evaluating temporal and spatial light exposure profiles for typical building occupants. In: Proceedings of the 11th annual symposium on simulation in architecture and urban design. Virtual event (Austria): Society for Computer Simulation International. p 55.

de Vries SW, Mardaljevic J. 2026. Code accompanying the paper: Performance verification of human field of view occluders for light measurement and simulation [script]. Version 1.0. 10.4121/a3b8c37a-de2d-48e4-9eee-ce456a2f98ce.

de Vries SW, Gkaintatzi-Masouti M, van Duijnhoven J, Mardaljevic J, Aarts MPJ. 2025. Recommendations for light-dosimetry field studies based on a meta-analysis of personal light levels of office workers. Lighting Res Technol. 57(1):47–70. 10.1177/14771535241248540.

de Vries SW, Mardaljevic J, van Duijnhoven J. 2026a. Impact of wear position on dosimeter performance: a hybrid measurement-simulation approach to quantify in-situ factors. npj Biol Timing Sleep. 3(1):20. 10.1038/s44323-026-00079-z.

de Vries SW, Mardaljevic J, van Duijnhoven J. 2026b. Impact of wear position on dosimeter performance: measurement validity under simulated indoor illumination. npj Biol Timing Sleep. 3(1):19. 10.1038/s44323-026-00073-5.

Dijk D, Archer SN. 2009. Light, sleep, and circadian rhythms: together again. PLOS Biol. 7(6):e1000145. 10.1371/journal.pbio.1000145.

Glickman G et al. 2003. Inferior retinal light exposure is more effective than superior retinal exposure in suppressing melatonin in humans. J Biol Rhythms. 18(1):71–79. 10.1177/0748730402239678.

Hartmeyer SL, Webler FS, Andersen M. 2023. Towards a framework for light-dosimetry studies: methodological considerations. Lighting Res Technol. 55(4-5):377–399. 10.1177/14771535221103258.

He S, Li H, Yan Y, Cai H. 2024. Capturing luminous flux entering human eyes with a camera, part 1: fundamentals. LEUKOS. 20(1):9–27. 10.1080/15502724.2022.2147942.

Illuminating Engineering Society. 1990. IES recommended standard file format for electronic transfer of photometric data. Journal of the Illuminating Engineering Society. 19(1):187–194. 10.1080/00994480.1990.10747955.

International Commission on Illumination. 2018. CIE S 026/E:2018 CIE system for metrology of optical radiation for ipRGC-influenced responses to light. CIE Central Bureau. 10.25039/S026.2018.

Knoop M et al. 2019. Methods to describe and measure lighting conditions in experiments on non-image-forming aspects. LEUKOS. 15(2-3):163–179. 10.1080/15502724.2018.1518716.

Krishnaswamy A, Baranosk GV, Rokne JG. 2004. Improving the reliability/cost ratio of goniophotometric comparisons. J Graph Tools. 9(3):1–20. 10.1080/10867651.2004.10504894.

Lasko TA, Kripke DF, Elliot JA. 1999. Melatonin suppression by illumination of upper and lower visual fields. J Biol Rhythms. 14(2):122–125. 10.1177/074873099129000506.

Lucas RJ et al. 2014. Measuring and using light in the melanopsin age. Trends Neurosci. 37(1):1–9. 10.1016/j.tins.2013.10.004.

Mardaljevic J. 1995. Validation of a lighting simulation program under real sky conditions. Lighting Res Technol. 27(4):181–188. 10.1177/14771535950270040701.

Mardaljevic J. 2001. The BRE-IDMP dataset: a new benchmark for the validation of illuminance prediction techniques. Lighting Res Technol. 33(2):117–134. 10.1177/136578280103300209.

Mardaljevic J. 2002. Quantification of parallax errors in sky simulator domes for clear sky conditions. Lighting Res Technol. 34(4):313–327. 10.1191/1365782802li055oa.

Mardaljevic J, Roy N. 2016. The sunlight beam index. Lighting Res Technol. 48(1):55–69. 10.1177/1477153515621486.

Mardaljevic J, Andersen M, Roy N, Christoffersen J. 2014. A framework for predicting the non-visual effects of daylight – part II: the simulation model. Lighting Res Technol. 46(4):388–406. 10.1177/1477153513491873.

Mardaljevic J, Cannon-Brookes S, Blades N, Lithgow K. 2021. Reconstruction of cumulative daylight illumination fields from high dynamic range imaging: theory, deployment and in-situ validation. Lighting Res Technol. 53(4):311–331. 10.1177/1477153520945755.

Moreno I, Sun C. 2008. LED array: where does far-field begin? Eighth International Conference on Solid State Lighting. International Society for Optics and Photonics. SPIE. p 70580R. 10.1117/12.795944.

Papamichael K, Klems J, Selkowitz SE. 1988. Determination and application of bidirectional solar-optical properties of fenestration systems. Lawrence Berkeley National Laboratory. LBL-25124.

Rea MS, Nagare R, Figueiro MG. 2021. Relative light sensitivities of four retinal hemifields for suppressing the synthesis of melatonin at night. Neurobiol Sleep Circadian Rhythms. 10:100066. 10.1016/j.nbscr.2021.100066.

Rüger M, Gordijn MCM, Beersma DGM, de Vries B, Daan S. 2005. Nasal versus temporal illumination of the human retina: effects on core body temperature, melatonin, and circadian phase. J Biol Rhythms. 20(1):60–70. 10.1177/0748730404270539.

Scartezzini JL, Compagnon R, Reecker C, Michel L. 1997. Bidirectional photogoniometer for advanced glazing materials based on digital imaging techniques. Int J Lighting Res Technol. 29(4):197–205. 10.1177/14771535970290040201.

Sliney DH. 1983. Eye protective techniques for bright light. Ophthalmology. 90(8):937–944. 10.1016/S0161-6420(83)80021-9.

Sliney D. 2019. Retinal exposure assessment – horizontal or vertical alpha irradiance or illuminance? In: Proceedings of the 29th CIE session. Washington D.C. (USA): CIE. 10.25039/x46.2019.OP22.

Smith JS, Kripke DF, Elliott JA, Youngstedt SD. 2002. Illumination of upper and middle visual fields produces equivalent suppression of melatonin in older volunteers. Chronobiol Int. 19(5):883–891. 10.1081/CBI-120014107.

Spitschan M et al. 2022. Verification, analytical validation and clinical validation (V3) of wearable dosimeters and light loggers. DIGITAL HEALTH. 8:20552076221144858. 10.1177/20552076221144858.

van Derlofske JF, Bierman A, Rea MS, Maliyagoda N. 2000. Design and optimization of a retinal exposure detector. Novel Optical Systems Design and Optimization III. International Society for Optics and Photonics. SPIE. p 60 – 70. 10.1117/12.402413.

van Duijnhoven J et al. 2025. Measuring light exposure in daily life: A review of wearable light loggers. Build Environ. 274:112771. 10.1016/j.buildenv.2025.112771.

Visser EK, Beersma DGM, Daan S. 1999. Melatonin suppression by light in humans is maximal when the nasal part of the retina is illuminated. J Biol Rhythms. 14(2):116–121. 10.1177/074873099129000498.

Ward GJ. 1992. Measuring and modeling anisotropic reflection. SIGGRAPH Comput. Graph. 26(2):265–272. 10.1145/142920.134078.

Ward GJ et al. 1998. Rendering with radiance: the art and science of lighting visualization. San Francisco: Morgan Kaufmann.

Watson AB, Yellott JI. 2012. A unified formula for light-adapted pupil size. J Vis. 12(10):12. 10.1167/12.10.12.

Xiao H, Cai H, Li X. 2021. Non-visual effects of indoor light environment on humans: A review. Physiol Behav. 228:113195. 10.1016/j.physbeh.2020.113195.

Zauner J, Broszio K, Bieske K. 2023. Influence of the Human Field of View on Visual and Non-Visual Quantities in Indoor Environments. Clocks Sleep. 5(3):476–498. 10.3390/clockssleep5030032.

